## Supplementary Information for "eQTL mapping in fetal-like pancreatic progenitor cells reveals early developmental insights into diabetes risk"

#### Supplemental Note 1: Characterization of iPSC-PPC fetal-like transcriptomes using scRNA-seq

Pancreatic progenitor cells (PPCs) are multipotent stem cells that have differentiated beyond the pancreatic foregut and can give rise to all pancreatic cell types (both endocrine and exocrine). PPCs are marked by co-expression of PDX1 and NKX6-1, which we quantified using flow cytometry (Figure 1B, Supplementary Figure 2-3, Supplementary Data 2). To characterize the cellular composition of iPSC-PPCs, we performed scRNA-seq on one iPSC clone (for PPC034) and ten iPSC-PPC samples (Supplementary Data 2). For four of the ten iPSC-PPC samples, we collected both freshly prepared (i.e., not cryopreserved) and cryopreserved cells to examine the effects of cryopreservation on expression quantification. For the remaining six iPSC-PPC samples, the cells were captured only as cryopreserved cells.

We detected eight distinct cell type clusters in iPSC-PPCs (Supplementary Data 2, Supplementary Data 3, Supplementary Data 4). To annotate each cluster, we compared the expression levels of pancreas development marker genes to those in the eight PPC cell types identified in a reference dataset (Supplementary Figure 4-6) [1]. The Veres et al. study [1] performed scRNA-seq over four stages of embryonic stem cell derived-PPC (ESC-PPC) differentiation and included cells from early onset of differentiation (pancreas induction) to more advanced stages (endocrine and exocrine induction), thus providing a valuable benchmark to characterize stem cell-derived PPC cellular heterogeneity. One of the eight clusters in our dataset expressed high levels of *POU5F1*, indicating that this cluster corresponded to the iPSC sample (Supplementary Figure 4-6, Supplementary Data 4). For the iPSC-PPC clusters, we identified: 1) early PPC, which expressed *GATA4*, *GATA6*, and *PDX1*, but not *NKX6-1* and corresponded to $PDX1^+$ progenitors in the Veres et al. study; 2) late PPC, which expressed both *PDX1* and *NKX6-1* and corresponded to $NKX6\text{-}1^+$ progenitors in the Veres et al. study; 3) endocrine cells, which expressed endocrine markers (*PAX6*, *CHGA*) and pancreatic hormones (*INS*, *GCG*, *SST*); and 4) non-endocrine cells, which expressed endothelial markers (*ESM1*, *FLT1*, *PLVAP*) but not mature ductal marker *KRT19* or mature acinar marker *PRSS1*, suggesting that these cells were likely ductal precursors (herein referred to as “early ductal”). We also identified a sub-population within late PPCs that expressed cell division markers (*TOP2A*, *CENPF*, and *AURKB*), indicating that these cells were replicating late PPC and corresponded to replicating $NKX6\text{-}1^+$ progenitors in the Veres et al. study (labeled as “replicating stage 4” in Supplementary Figure 4). Of note, we identified two cell types from very early pancreas development that were not captured in the Veres et al. dataset, one of which represented mesendoderm (*COL1A1/2*) and the other early definitive endoderm (“early DE”; *AFP*, *APOA2*).

We next examined the extent to which iPSC-PPC cellular heterogeneity was reflected in the flow cytometry analysis. We found that the percentage of late PPCs (including replicating cells) in scRNA-seq was significantly correlated with the percentage of $PDX1^+/NKX6\text{-}1^+$ cells measured by flow cytometry ($R = 0.857$, $p = 0.00154$, Pearson’s correlation, Supplementary Figure 7A, Supplementary Data 3). We also deconvoluted cell type proportions in bulk RNA-seq samples of iPSC-PPC using cell type-specific markers in scRNA-seq and found that the estimated proportions for late PPC highly corresponded with flow cytometry measurements of $PDX1^+/NKX6\text{-}1^+$ cells ($R = 0.702$, $p < 2.2\times10^{-16}$, Pearson’s correlation, Supplementary Figure 7B, Supplementary Data 5). These results show that the flow cytometry analysis accurately captured the fraction of late PPCs in the 107 iPSC-PPCs.

We next asked whether cryopreservation affected gene expression profiling in iPSC-PPC. Because gene expression differences between samples are largely driven by cellular heterogeneity, we sought to compare the cell type proportions in scRNA-seq between cryopreserved cells and freshly prepared cells for the four iPSC-PPC samples sequenced with both preparations. We observed that the cellular proportions between both preparations for late PPC, early PPC, mesendoderm, and endocrine were significantly correlated (Supplementary Figure 8, Supplementary Data 3). These results show that cryopreservation did not impact gene expression levels.

#### Supplemental Note 2: Analysis of iPSC-PPC $e_{AS}$QTLs

##### 2a. COLOC analysis identifies shared $e_{AS}$QTLs between the three pancreatic tissues

Similar to the $e_g$QTL analysis, we performed colocalization between nearby pairs of iPSC-PPC $e_i$QTLs, adult islet exon eQTLs, and adult whole pancreas splicing eQTLs. Hereafter, we refer to these three different types of eQTLs as $e_{AS}$QTLs given their functional properties related to alternative splicing [2–4] (Figure 1E). We considered only $e_{AS}$QTLs that had at least one variant with causal PP ≥ 1%, were outside the MHC region, and associated with genes annotated in GENCODE version 34 [5], therefore retaining 3,959 iPSC-PPC, 4,939 adult islets, and 2,077 adult whole pancreas $e_{AS}$QTLs. From colocalization, we identified a total of 4,868 pairs of $e_{AS}$QTLs that displayed high evidence of colocalization with PP.H4 ≥ 80% (Supplementary Data 11). We also observed that $e_{AS}$QTLs for the same gene may colocalize with each other indicating that a single causal variant may impact multiple splicing processes for a gene.

We next identified tissue-unique singleton and combinatorial $e_{AS}$QTLs. Using the same approach in the $e_g$QTL analysis, we identified 631 iPSC-PPC, 1,522 adult islets, and 431 adult whole pancreas singleton $e_{AS}$QTLs (i.e., $e_{AS}$QTLs that neither colocalized nor were in LD ($r^2 < 0.2$ within 500 Kb or outside of 500 Kb if LD not available) with nearby $e_{AS}$QTLs, indicating that their underlying causal variants affect alternative splicing of a single transcript specifically during early pancreas development or in one of the adult pancreatic tissues (Supplementary Figure 11A, Supplementary Data 11, Supplementary Data 12). To identify tissue-unique combinatorial $e_{AS}$QTLs, we created a network using the 4,868 pairwise colocalizations and then filtered modules that failed specific module and LD criteria (see Methods). We identified 980 $e_{AS}$QTL modules in total, averaging ~3 $e_{AS}$QTLs per module (range: 2-13) (Supplementary Data 12, Supplementary Data 13). 344 (35.1% of 980) $e_{AS}$QTL modules were tissue-unique, of which 124 were fetal-like iPSC-PPC-unique, 203 adult islet-unique, and 17 adult whole pancreas-unique, and comprised 266, 452, 37 $e_{AS}$QTLs, respectively (Supplementary Figure 11B). The remaining 636 (64.9% of 980) $e_{AS}$QTL modules were shared between multiple pancreatic tissues, 225 of which were shared between only the two adult pancreatic tissues ("adult-shared"), 139 shared between iPSC-PPC and adult islets ("fetal-islet"), 58 shared between only iPSC-PPC and adult whole pancreas ("fetal-whole-pancreas"), and 214 shared between all three pancreatic tissues ("fetal-adult") (Supplementary Figure 11B). Together, the 411 (139 + 58 + 214) modules shared between iPSC-PPC and an adult pancreatic tissue comprised 802 iPSC-PPC, 561 adult islets, and 318 adult whole pancreas $e_{AS}$QTLs (Supplementary Data 12, Supplementary Data 13).

Altogether, from this analysis, we identified 897 (22.7% of 3,959) iPSC-PPC-unique $e_{AS}$QTLs, of which 631 (70.3%) functioned as singletons and 266 (29.7%) in modules, while 802 (20.3% of 3,959) were shared with at least one adult pancreatic tissue (Supplementary Data 12, Supplementary Data 13). The remaining 2,260 (57.1% of 3,959) iPSC-PPC $e_{AS}$QTLs failed either module or LD criteria (see Methods). In adult whole pancreas, we observed much fewer tissue-unique

$e_{AS}$QTL modules compared to $e_g$QTLs because each gene only had one significant sQTL and therefore unlikely to colocalize with one another. On the other hand, in iPSC-PPCs and adult islets, genetic variants affect the expression of multiple isoforms or exons corresponding to the same gene in the same tissue; and hence, compared with the $e_g$QTLs, a greater fraction of the iPSC-PPC-unique and the adult islet-unique $e_{AS}$QTLs were combinatorial. Similar to the $e_g$QTLs, the vast majority of tissue-unique regulatory variants were singletons, potentially due to large differences in alternative splicing between fetal and adult pancreatic tissues, which is known to exist for other tissue-types [6–8]. Additionally, there may be other distinct properties $e_i$QTLs, exon eQTLs, and splicing eQTLs that we did not account for.

**2b. Characterization of fetal-adult-shared $e_{AS}$QTL modules in iPSC-PPC**

We next determined the fraction of shared genetic loci associated with alternative splicing of the same or different genes between fetal-like iPSC-PPC and the two adult pancreatic tissues. Similar to the $e_g$QTL analysis, we focused on the 411 $e_{AS}$QTL modules with both iPSC-PPC and adult $e_{AS}$QTLs ("fetal-adult", "fetal-islet", "fetal-whole-pancreas") and compared the genes associated with each module. We identified: A) 149 modules that were associated with the same gene between fetal-like iPSC-PPC and only one of the two adult pancreatic tissues (1 gene per module); B) 85 modules associated with the same gene in all three tissues (1 gene per module); C) 93 modules associated with 2-5 genes, of which some genes were shared but at least one gene was different between iPSC-PPC and at least one adult tissue; D) 57 modules associated with different genes between iPSC-PPC and only one of the two adult tissues (2-5 genes per module); E) the remaining 27 modules associated with different genes between fetal-like and both the two adult tissues (range: 2-5 genes per module) (i.e., there is no overlap of genes between the two developmental stages; Supplementary Figure 11C, Supplementary Data 13). 43.1% (93 + 57 + 27 = 177 / 411; categories C-E) of the fetal-adult-shared modules displayed functional plasticity, in which the underlying regulatory variants were associated with splicing events for multiple different genes. These modules comprised 386 iPSC-PPC, 361 adult islets, and 184 adult whole pancreas $e_{AS}$QTLs (Supplementary Data 12, Supplementary Data 13).

#### Supplementary Figure 1: Subject and sample characteristics of iPSC-PPC cohort

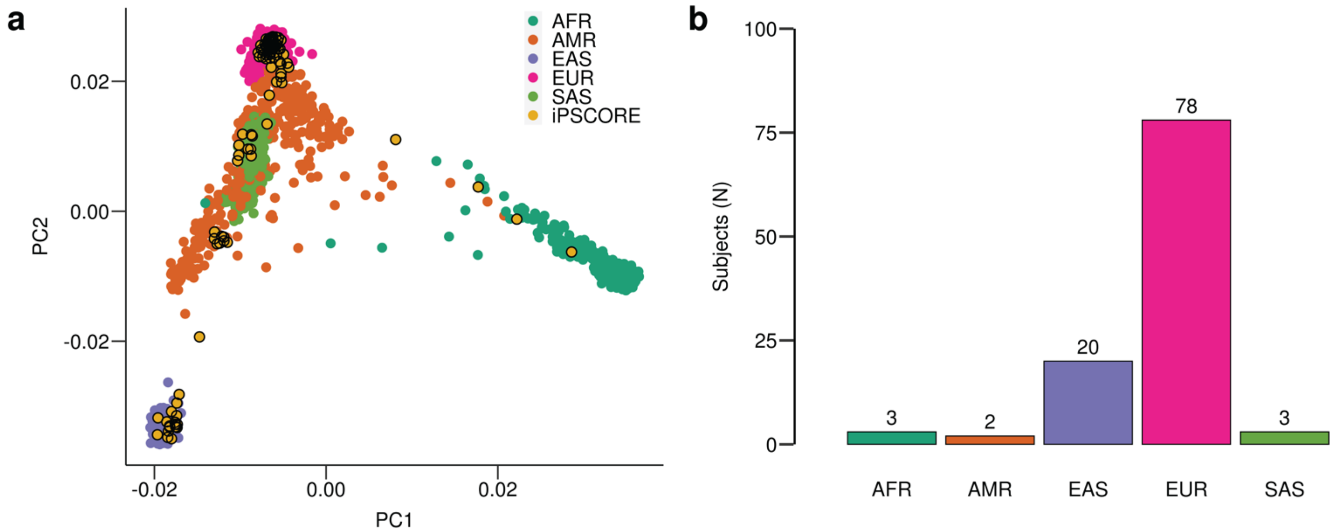


(**a**) Scatter plot showing the first two genotype principal components of the 106 iPSCORE subjects in relation to the 2,504 subjects in the 1000 Genomes Project (1KGP) Phase 3 dataset. Color indicates the five 1KGP superpopulations (AFR, AMR, EAS, EUR, and SAS). Black bordered yellow circles correspond to the 106 iPSCORE individuals in this study (Supplementary Data 1). (**b**) Bar plot showing the distribution of the 106 iPSCORE subjects across different 1KGP superpopulations. Subjects were assigned to the most similar 1KGP population in a previous study using linear discriminant analysis [9].

#### Supplementary Figure 2: Measurement of $PDX1^{+}$ and $NKX6\text{-}1^{+}$ by flow cytometry for 107 iPSC-PPC samples

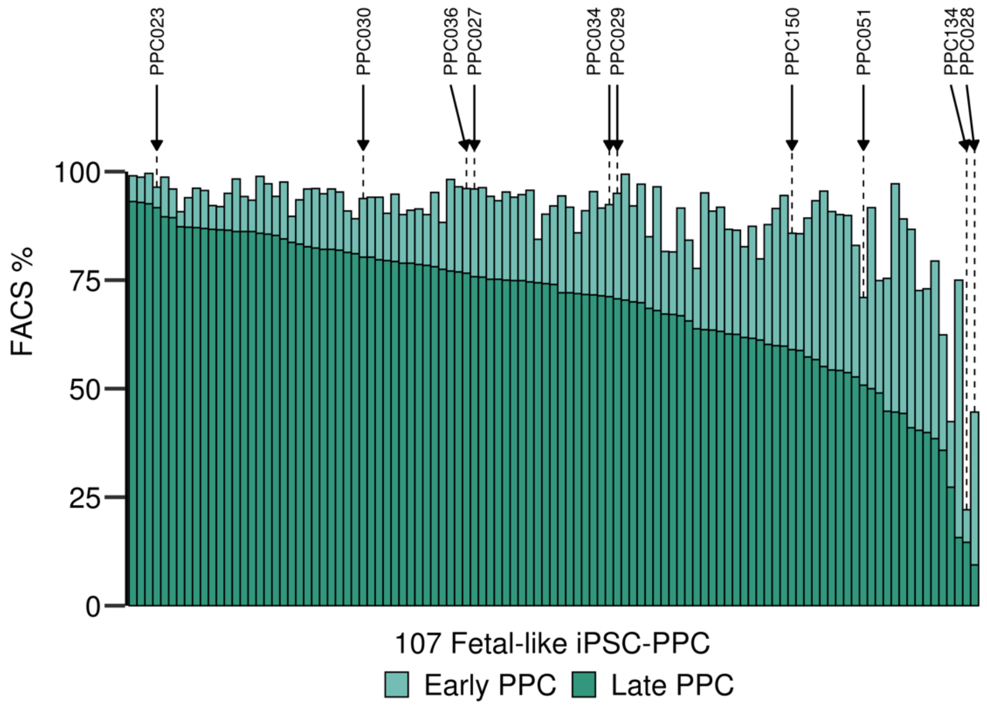


Bar plot showing the percentage of late PPC and early PPC cells detected by FACs for all of the 107 differentiations (Supplementary Data 2). The ten highlight samples by arrows indicate samples used for scRNA-seq examination. We show that the samples selected for scRNA-seq varied in late PPC percentage. The two iPSC-PPC samples derived from the same iPSC line (PPC029 and PPC036) had similar percentages of late PPCs (70.7% and 76.6%, respectively).

**Supplementary Figure 3: Flow cytometry results for the ten iPSC-PPC samples used in single-cell analysis**

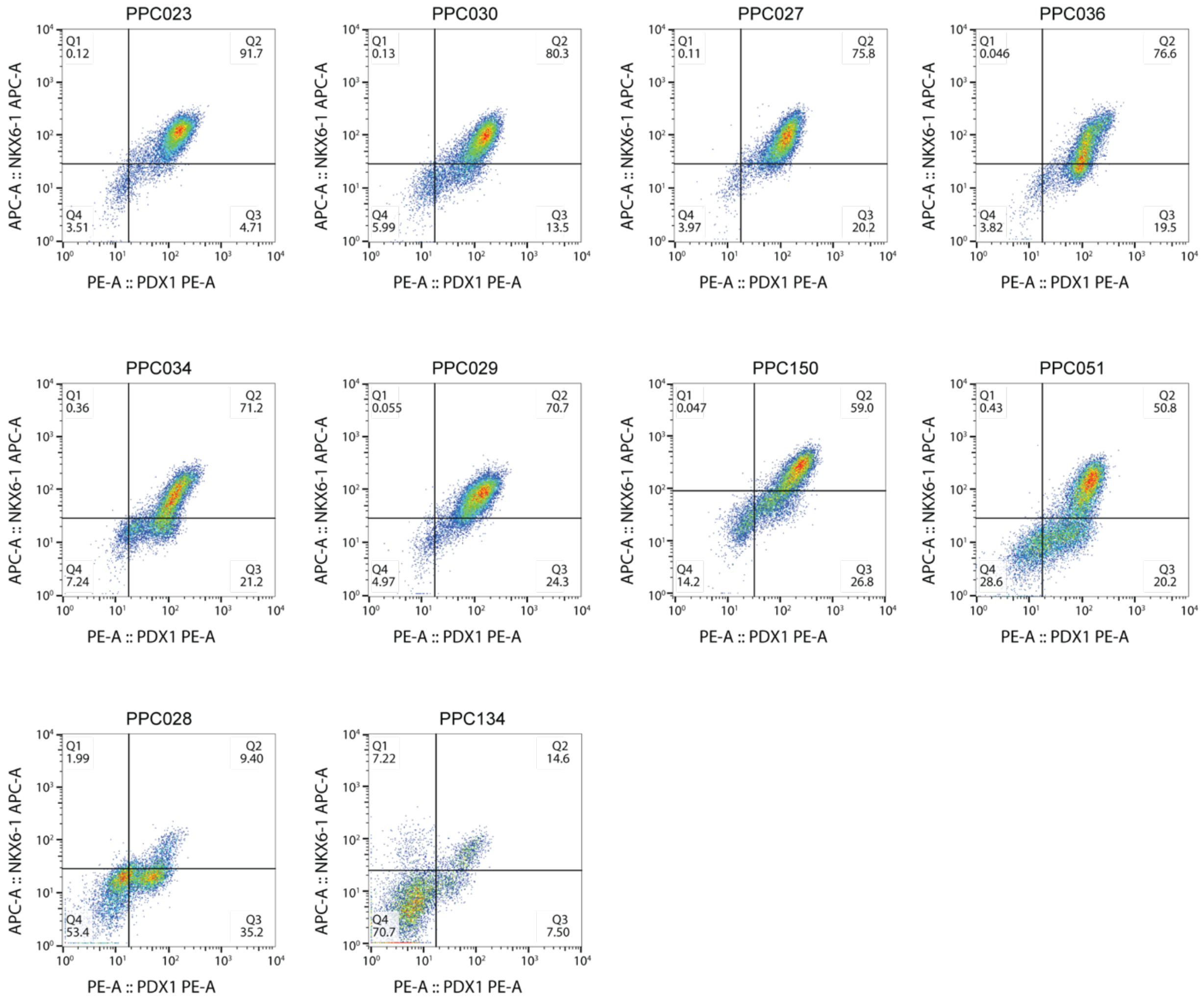


Flow cytometry analysis at D15 of the ten iPSC-PPC samples that underwent scRNA-seq (Supplementary Figure 2). The percentage of cells stained for PDX1 (X-axis) and NKX6-1 (Y-axis) were measured. Differentiations PPC029 and PPC036 were from the same iPSC line and showed similar percentages of double-positive cells. Measurements for all 107 iPSC-PPC differentiations are provided in Supplementary Data 2 and shown in Supplementary Figure 2.

#### Supplementary Figure 4: Single cell characterization of fetal-like iPSC-PPC

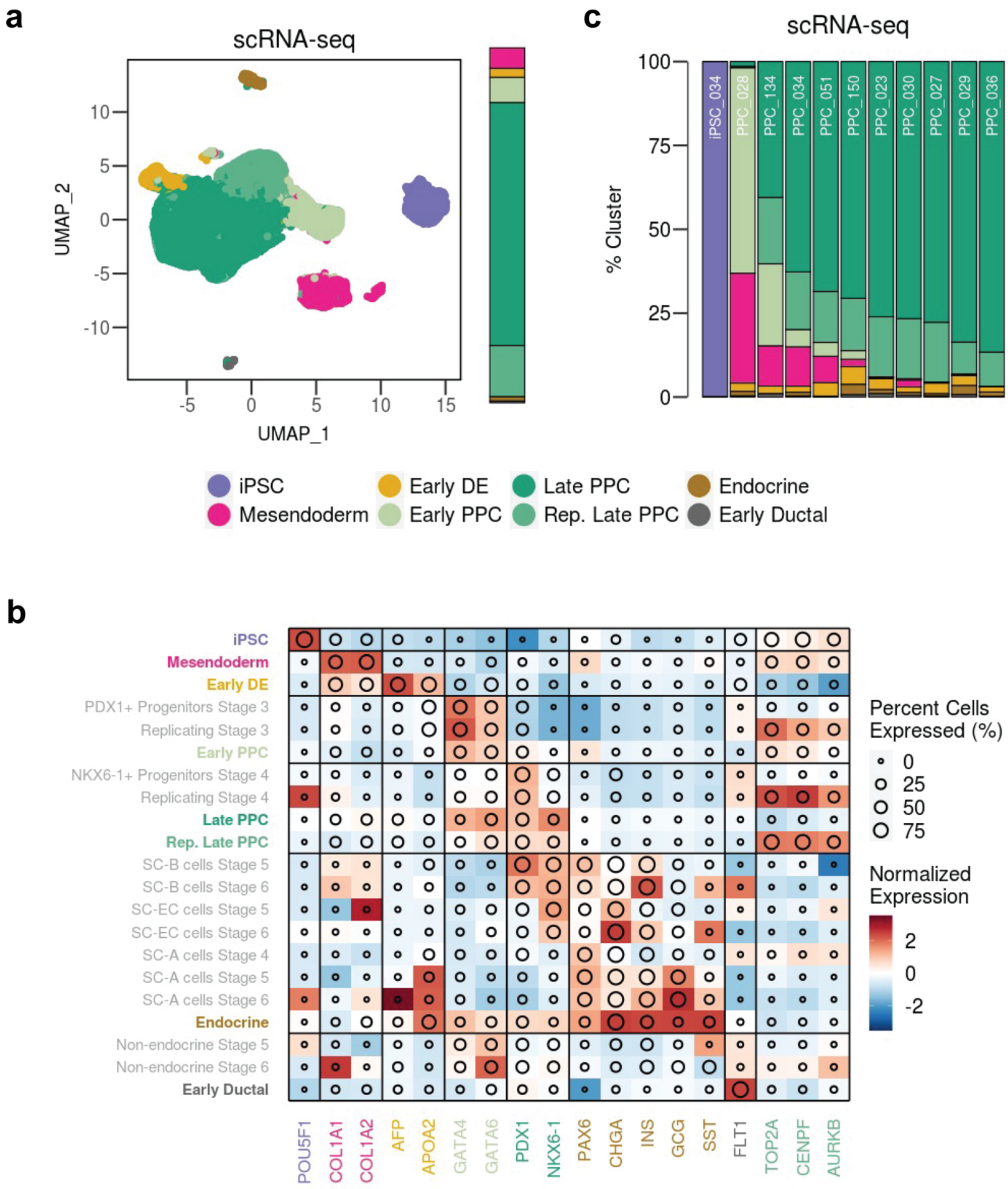


We characterized the cellular composition of iPSC-PPC using scRNA-seq of one iPSC (for PPC034 differentiation) and ten iPSC-PPC samples with variable percentages of double-positive cells (range: 9.4% - 91.7%) (Supplementary Figure 2, Supplementary Figure 3). We identified eight distinct cell populations, corresponding to iPSC (*POU5F1*), mesendoderm (*COL1A1/2*), early definitive endoderm (early DE; *AFP*, *APOA2*), early PPC (*GATA, GATA6, PDX1*), late PPC (*PDX1* and *NKX6-1*), replicating late PPC (*PDX1, NKX6-1, TOP2A, CENPF, AURKB*), endocrine (*PAX6, CHGA, INS, GCG, SST*), and early ductal (*FLT1*) (Supplementary Figure 4C, Supplementary Data 3). We observed highly similar gene expression profiles between the cell types identified in iPSC-PPC and those identified in an ESC-derived PPC (ESC-PPC) reference dataset [1].

(**a**) UMAP plot of scRNA-seq data from 84,225 single cells from one iPSC and ten iPSC-PPC samples. Each point represents a single cell color-coded by its assigned cluster. To the right of the UMAP plot, we show relative proportion of cells associated with each cell type (iPSC cells excluded). We show that the vast majority of cells in iPSC-PPCs were late PPCs. (**b**) Heatmap comparing the Z-normalized expression of known marker genes between iPSC-PPC and cells from the reference ESC-PPC study [1]. Color intensity indicates the mean Z-normalized expression across all cell types, and the diameter indicates the percentage of cells expressing the markers above the threshold of 1% of the maximum expression value. Clusters labeled in color correspond to the iPSC-PPC clusters. Clusters labeled in grey correspond to ESC-PPC clusters [1]. (**c**) Stacked bar plot showing the relative proportion of cells from each sample assigned to each cluster in scRNA-seq. Color-coding corresponds to the clusters in panel **a**. Samples with the least number of late PPC cells correspond to those with weaker differentiation efficiency based on FACS, and contain more cells of primitive state compared to the other samples.

### Supplementary Figure 5: Clustering of iPSC-PPC scRNA-seq at two additional resolutions

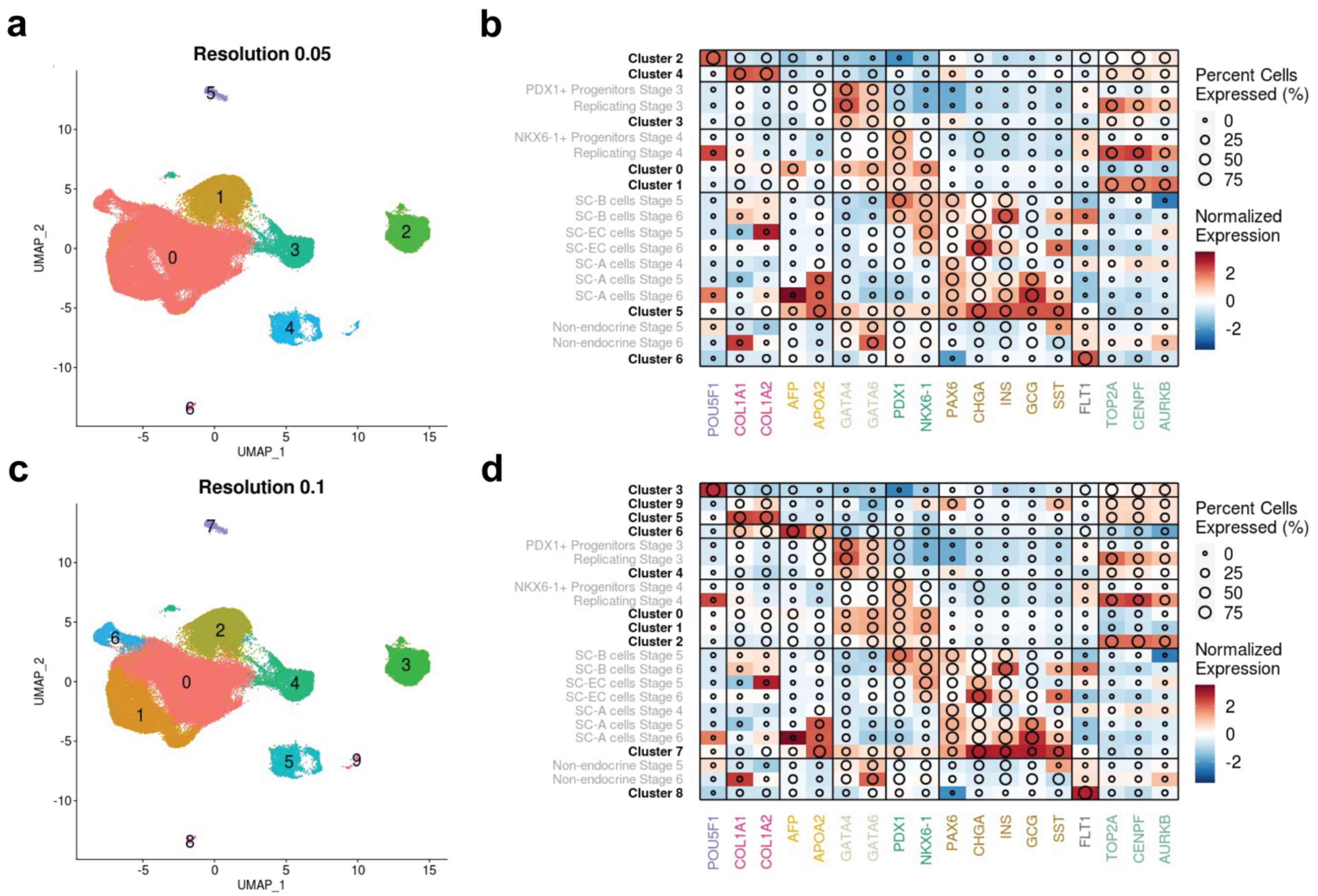


As described in the Methods, we performed clustering at three different resolutions: 0.05, 0.08 (shown in Supplementary Figure 4A), and 0.1, to annotate cell populations in the fetal-like iPSC-PPCs. Compared with resolution 0.08, resolution 0.05 (panel **a, b**) collapsed both early DE and late PPC as cluster 0, while resolution 0.1 (panel **c**) further divided late PPCs into two subclusters (cluster 0 and cluster 1). For resolution 0.1, we examined the expression profiles of the two subclusters in late PPCs and found that cluster 1 expressed similar levels of *PDX1* and *NKX6-1* as cluster 0 (panel **d**). Therefore, we used resolution 0.08 for downstream analyses (Supplementary Figure 4A). Color intensity indicates the mean Z-normalized expression across all cell types, and the diameter indicates the percentage of cells expressing the markers above the threshold of 1% of the maximum expression value. Cluster labels in black correspond to the iPSC-PPC clusters in the UMAP plots on the left. Cluster labels in grey correspond to ESC-PPC clusters [1].

**Supplementary Figure 6: Expression of 18 marker genes in each of the eight scRNA-seq clusters**

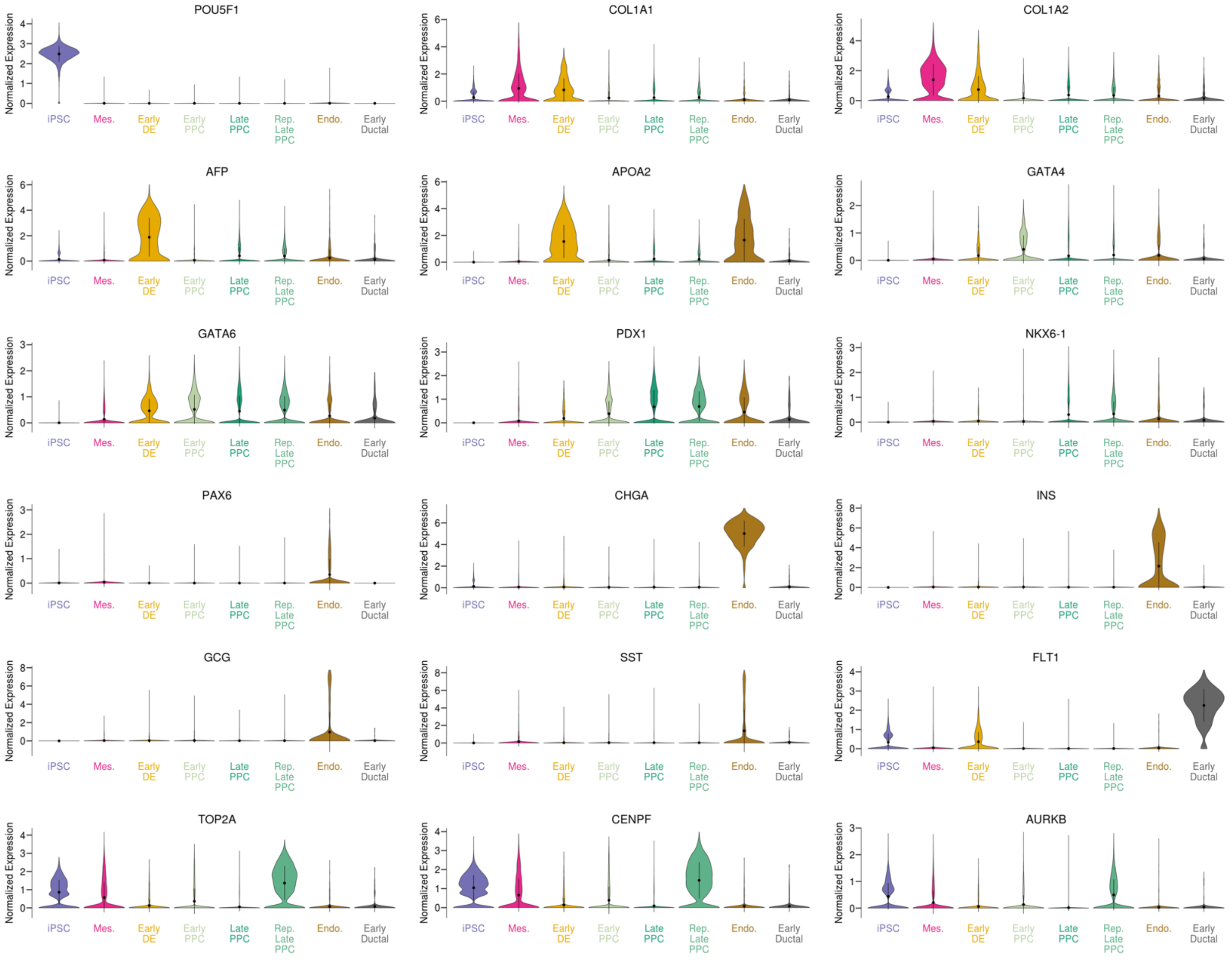


Violin plots showing the distribution of normalized expression for marker genes described in Supplementary Figure 4B for each iPSC-PPC scRNA-seq cluster in Supplementary Figure 4A. Center points in each violin represents the mean expression value. Error bars represent the standard deviation of expression across the cells.

#### Supplementary Figure 7: Correlation between flow cytometry and cell type proportions

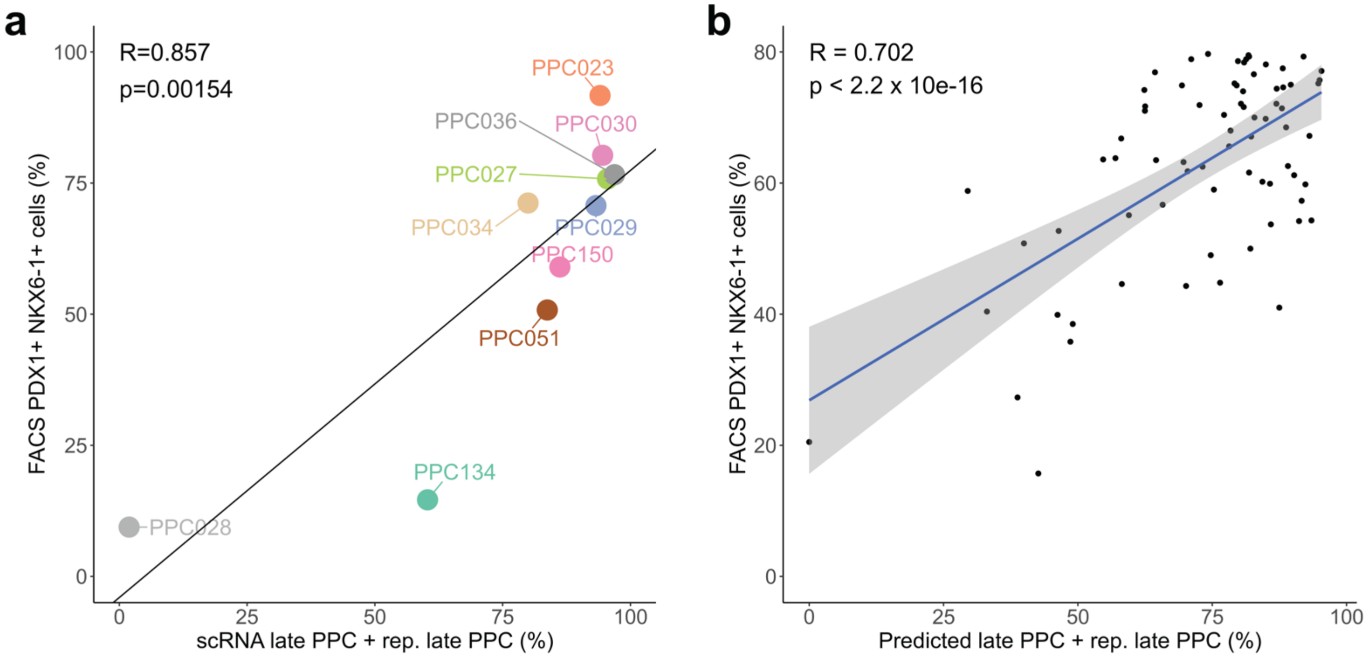


(**a**) To determine the correspondence between flow cytometry and scRNA-seq, we compared the percentage of double-positive cells in each iPSC-PPC samples as measured by flow cytometry (Y-axis; Supplementary Data 2, Supplementary Figure 2, Supplementary Figure 3) to the percentage of cells annotated as late PPC or replicating late PPC in scRNA-seq (X-axis; Supplementary Data 3, Supplementary Figure 4C). We found that independent measurements from flow cytometry and scRNA-seq were positively correlated (R = 0.857, p = 0.00154, spearman correlation). (**b**) We next examined the correspondence between the percentage of late PPCs from FACs (PDX1+/NKX6-1+) (Y-axis; Supplementary Data 2, Supplementary Figure 2, Supplementary Figure 3) and the estimated percentage of late PPCs (including replicating late PPCs) in bulk RNA-seq using CIBERSORTx [10] (X-axis; Supplementary Data 5). As expected, we observed a strong correlation between the two measurements ($R = 0.702$, $p < 2.2 \times 10^{-16}$), indicating that the cellular heterogeneity captured by flow cytometry is represented in the iPSC-PPC bulk expression profiles.

#### Supplementary Figure 8: Fresh and cryopreserved cells in scRNA-seq

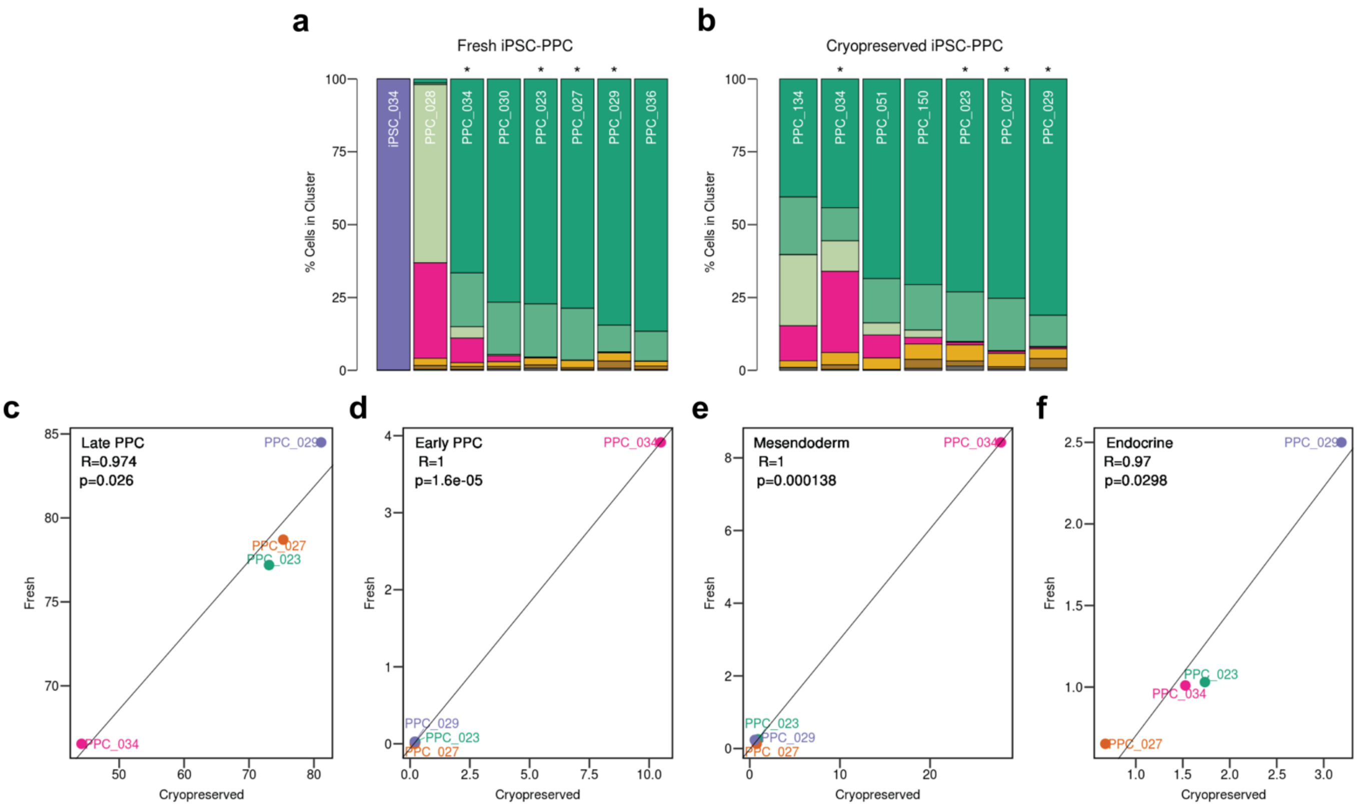


To determine if cryopreservation affects gene expression in iPSC-PPC, we asked whether the cell type proportions differed between matched fresh and cryopreserved samples for four iPSC-PPCs (PPC034, PPC023, PPC027, and PPC029; indicated by an asterisk in panel **a** and **b**). We compared the relative proportion of cells in early PPC, late PPC, mesendoderm, and endocrine clusters, and observed that fresh and cryopreserved samples were highly correlated. These results suggest that cell cryopreservation does not affect relative gene expression between samples, and therefore, can be used to characterize the cellular heterogeneity of iPSC-PPC. The Supplementary Figure hows: (**a**) Stacked bar plots showing the percentage of cells in each of the eight freshly prepared samples (one iPSC and seven iPSC-PPC) according to their cell type using the same color coding as Supplementary Figure 4A. Asterisks indicate the four iPSC-PPC samples with matched cryopreserved preparations. (**b**) Stacked bar plots showing the percentage of cells in each of the seven cryopreserved samples (all iPSC-PPC) according to their cell type using the same color coding as Supplementary Figure 4A. (**c-f**) For the four iPSC-PPC samples with matched fresh and cryopreserved samples, we show the association between each preparation by comparing the percentage of cells for early PPC (**c**), late PPC (**d**), mesendoderm (**e**), and endocrine (**f**).

#### Supplementary Figure 9: iPSC-PPCs represent a fetal-like state of the pancreas

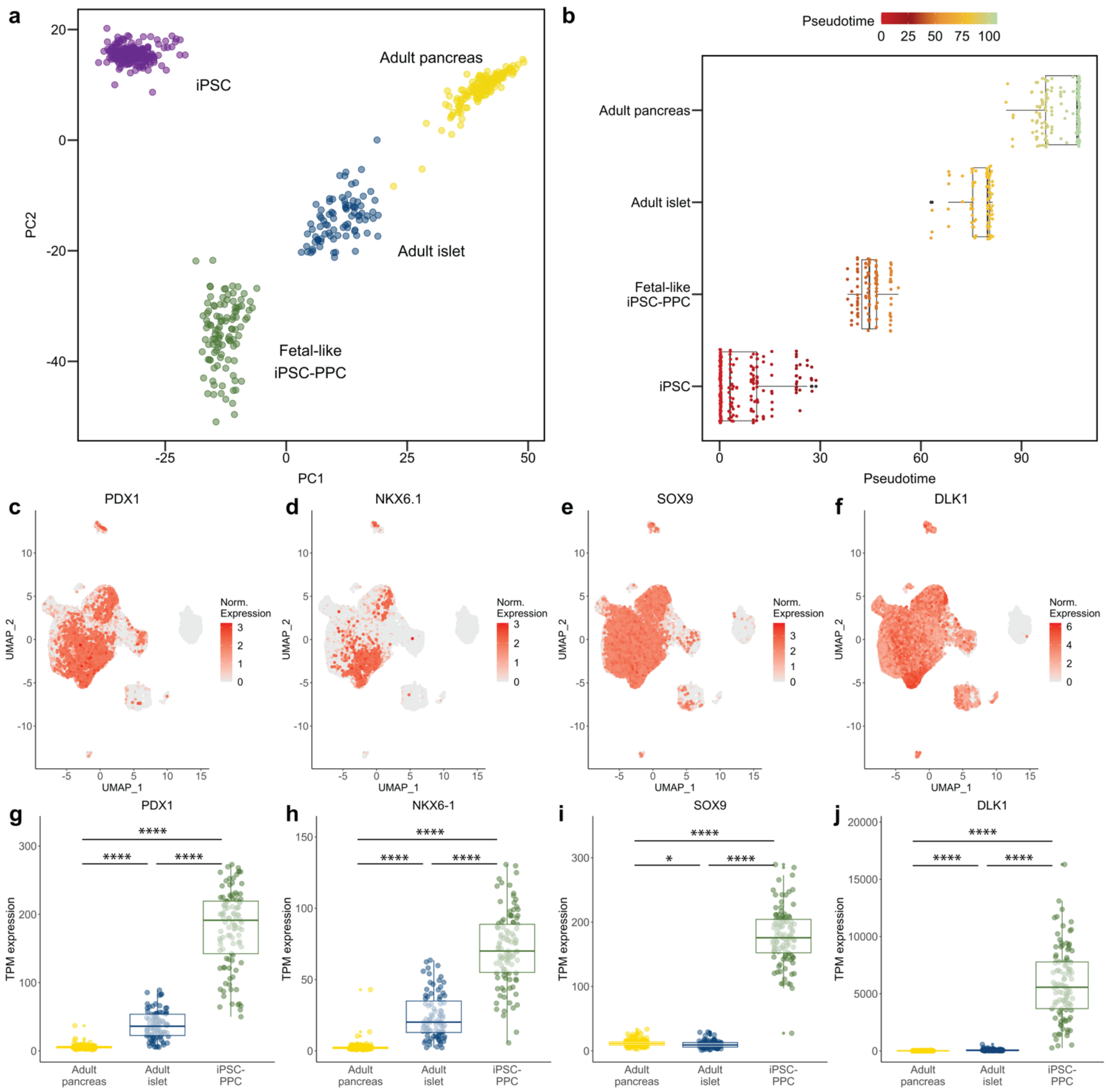


(**a**) Scatter plot showing the PCA distributions of the bulk transcriptomes of 213 iPSCs [11], 107 fetal-like iPSC-PPCs, 87 adult islets [12], and 176 adult whole pancreas [13] using the top most variable genes across all samples in bulk RNA-seq (Supplementary Data 6). (**b**) Box plot showing the pseudotime for each of the four tissues (213 iPSCs, 107 fetal-like iPSC-PPCs, 87 adult islets, and 176 adult whole pancreas) presented as +/- 1.5 interquartile range with the median line denoted at the center. Pseudotime was estimated using Monocle [14] where time was rooted at 0 in iPSCs. (**c-f)** UMAP plots showing the expression of fetal pancreatic genes [15–21] (*PDX1*, *NKX6-1*, *SOX9*, and *DLK1*) in early and late PPCs in iPSC-PPC (cell types labeled in Supplementary Figure 4A). (**g-j)** Box plots showing the expression of the same fetal pancreatic genes in panels **c-f** in bulk RNA-seq of 107 iPSC-PPCs (green), 87 adult islets (blue), and 176 adult whole pancreas (yellow). Paired student T-test was performed to evaluate the significance of expression differences between tissues. * = p < 0.05, ** = p < 0.01, *** = p < 0.001, **** = p < 0.0001. Box plots indicate median interquartile range (IQR), and 1.5 x IQR. These results confirm that iPSC-PPCs represent an early developmental time point compared to adult human islets and whole pancreas.

#### Supplementary Figure 10: Functional enrichment of overlapping $e_g$QTLs and $e_i$QTLs

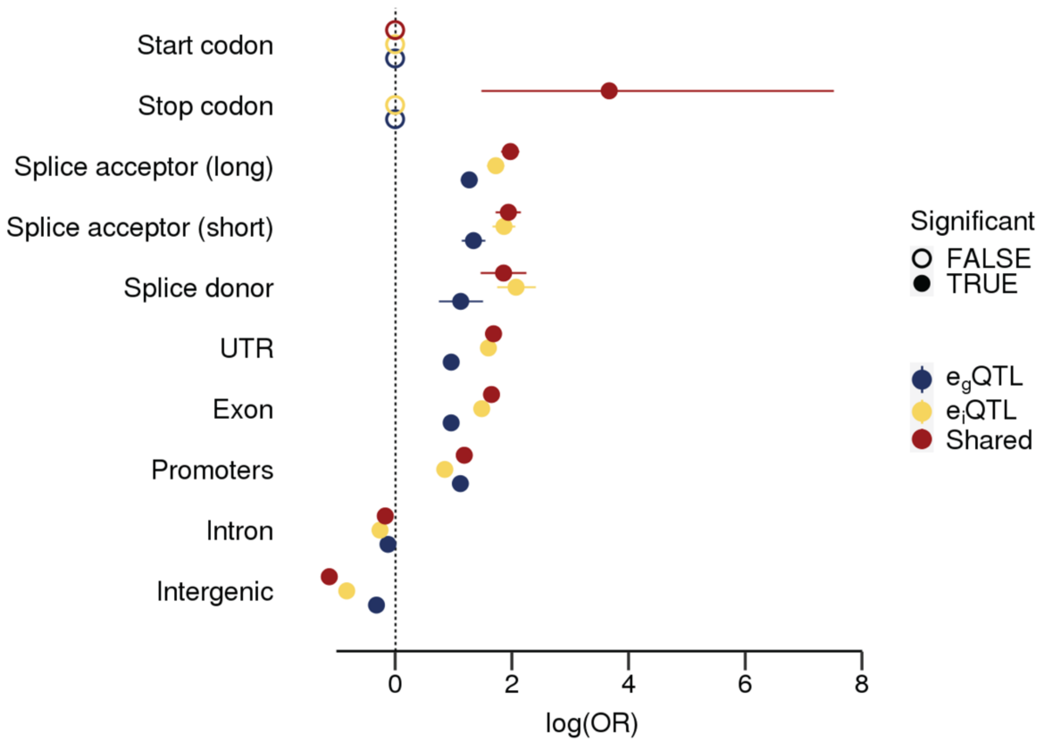


We divided the eQTL signals into three groups: 1) $e_g$QTLs that did not colocalize with $e_i$QTLs, 2) $e_i$QTLs that did not colocalize with $e_g$QTLs, and 3) eQTLs that had colocalized $e_g$QTLs and $e_i$QTLs ("shared", PP.H4 ≥ 80%). For non-colocalized $e_g$QTL and $e_i$QTL signals, we obtained all SNPs with causal PP ≥ 5% (Supplementary Data 8, Supplementary Data 11). For colocalized eQTLs, we used the predicted causal SNPs underlying both associations (output from colocalization). We intersected the SNP overlap with genomic regions (Y-axis) and performed a two-sided Fisher's Exact Test to calculate the enrichment of each eQTL group against a null background set of 20,000 variants. We found that overlapping eQTL signals were enriched in both regulatory and splice sites while non-overlapping $e_g$QTL and $e_i$QTLs were more enriched in their respective regions against each other. For example, $e_g$QTLs displayed a stronger enrichment for promoter regions compared to $e_i$QTLs while $e_i$QTLs were more enriched in splice sites compared to $e_g$QTLs, consistent with Figure 1E and other studies [2–4]. P-values were Benjamini-Hochberg-corrected and considered significant if the corrected p-values < 0.05. Non-significant results were set to log(odds ratio) = 0. Error bars represent 95% confidence intervals for the odds ratios.

#### Supplementary Figure 11: Singleton and combinatorial $e_{AS}$QTLs

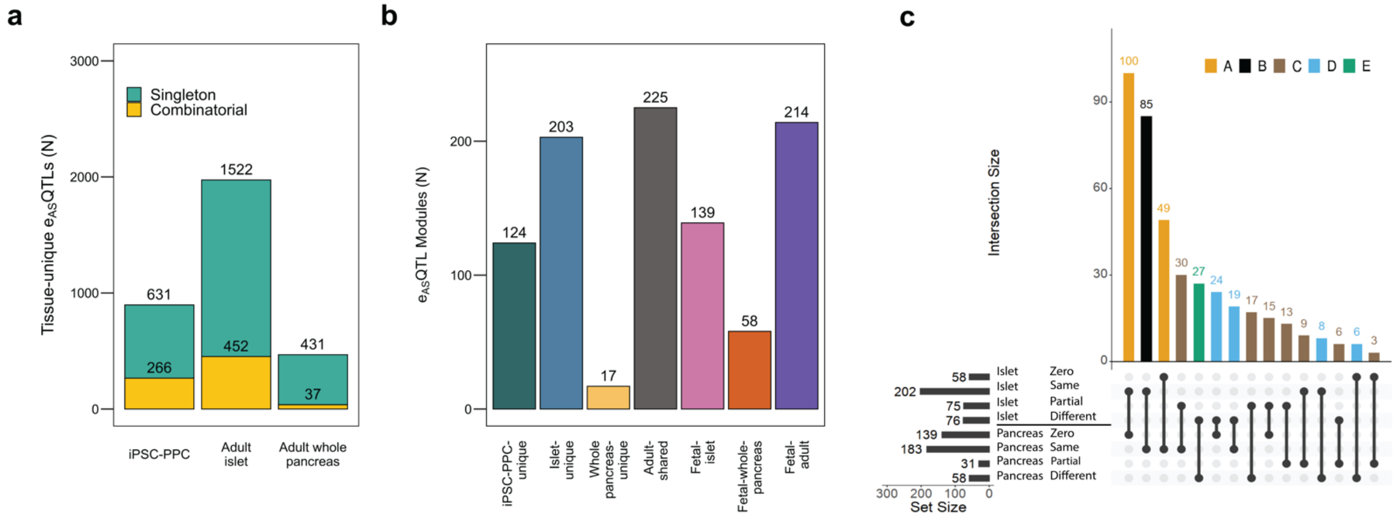


(**a**) Bar plot showing the number of $e_{AS}$QTLs that were classified as tissue-unique singletons (green) or in combinatorial (yellow) associations in each of the three pancreatic tissues. (**b**) Bar plot showing the number of $e_{AS}$QTL modules identified for each annotation. See Results and Methods for the description of each module category. (**c**) Number of $e_{AS}$QTL modules based on eGene overlap between iPSC-PPC and the two adult pancreatic tissues (Supplementary Data 13). "Zero" indicates that the module does not contain an $e_{AS}$QTL in the respective adult tissue. "Same" indicates that the module contains only $e_{AS}$QTLs corresponding to the same eGenes in iPSC-PPC and the adult tissue. "Partial" indicates that the module contains $e_{AS}$QTLs corresponding to partially overlapping eGenes between iPSC-PPC and the adult tissue. "Different" indicates that the module contains only $e_{AS}$QTLs corresponding to different eGenes between iPSC-PPC and the adult tissue.

#### Supplementary Figure 12: Definition of a GWAS locus

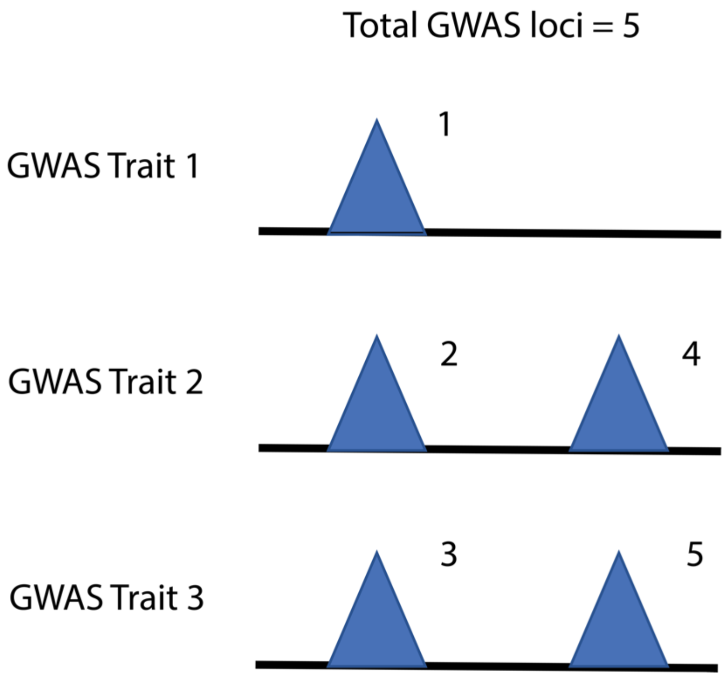


Given that some traits are highly correlated with one another, we observed some eQTLs that colocalized with GWAS variants associated with more than one trait. We considered each combination of colocalized eQTL-GWAS trait variants as a separate GWAS locus. The example in the Supplementary Figure hows two eQTL signals that overlap with either three or two traits, which we count as five GWAS loci. Considering single eQTLs we observed 183 GWAS loci and considering combinatorial eQTLs we observed 129 GWAS loci (Supplementary Figure 13).

#### Supplementary Figure 13: Singleton and module colocalization with pancreatic traits and disease

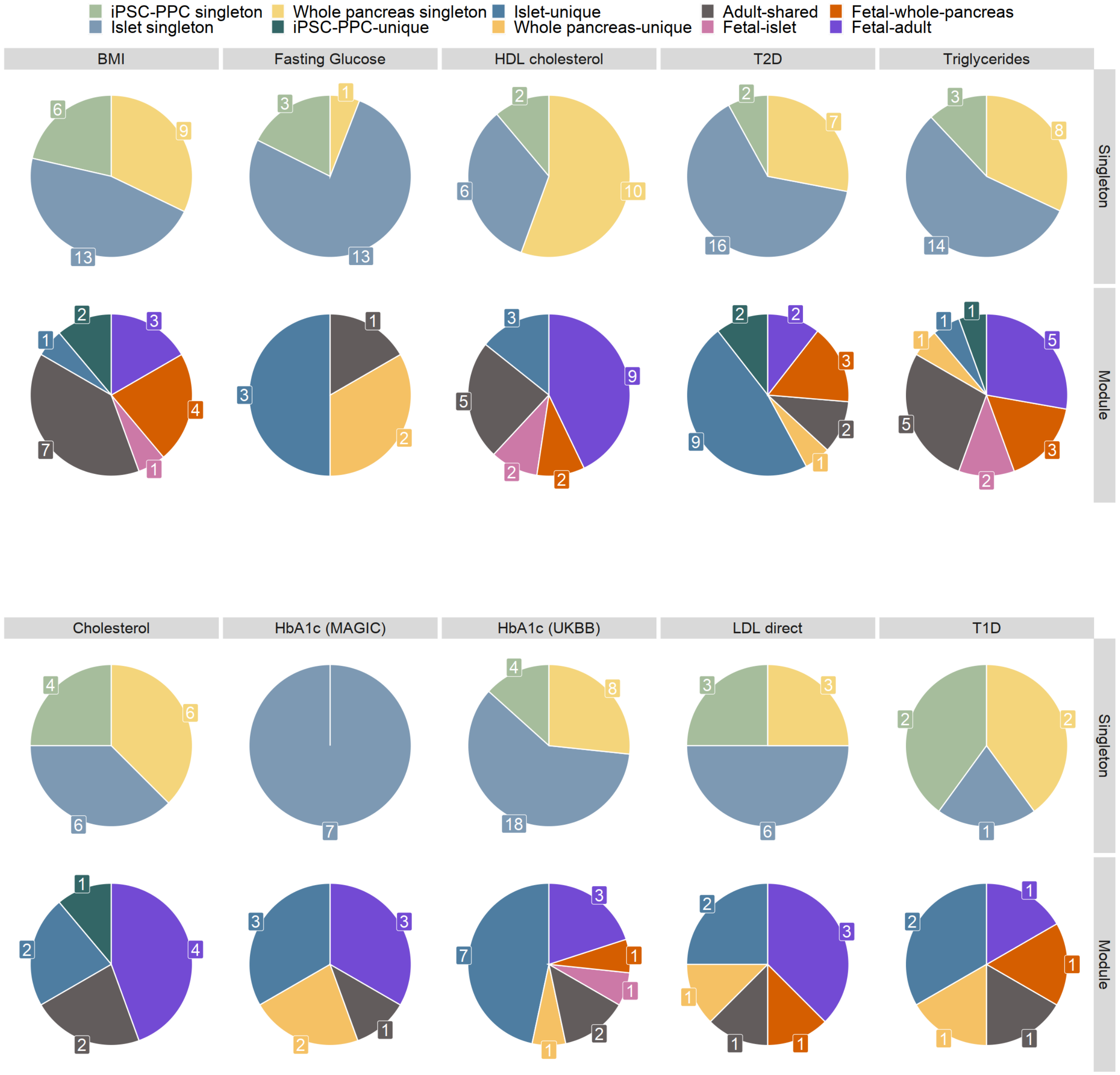


Pie charts summarizing number of colocalizations between eQTLs and GWAS variants for each of the ten traits. Labels indicate the number of singleton eQTLs (top rows) and modules (bottom rows) that colocalized with a GWAS signal. For example, for T1D, two iPSC-PPC-unique singleton (light green), one adult islet-unique singleton (light blue), and two adult whole pancreas-unique singleton (light yellow) eQTLs colocalized with a T1D-risk signal. Similarly, we identified two adult islet-unique eQTL modules (blue), one fetal-adult eQTL modules (orange), and etc., that colocalized with T1D-risk variants.

### Supplementary Figure 14: $e_gQTL$ associations for GWAS-associated eGenes in iPSC-PPC

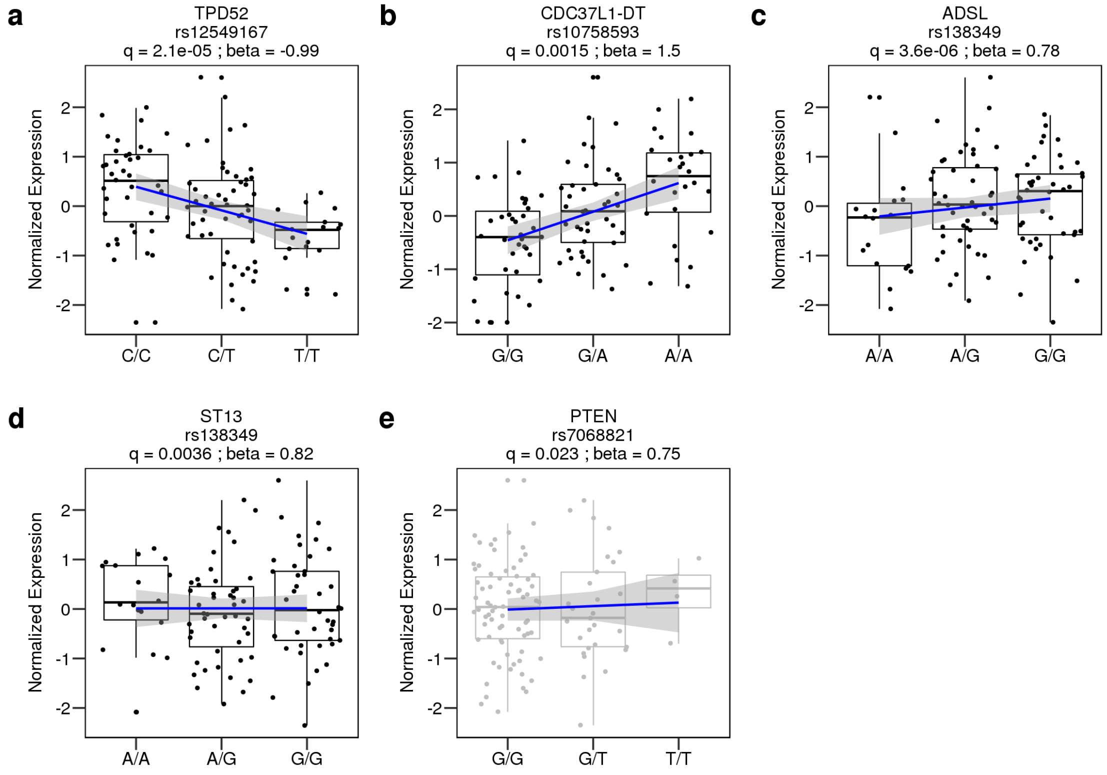


For each candidate susceptibility eGene, we show the association between genotype of the lead candidate causal variant (predicted from GWAS-eQTL colocalization) and their normalized gene expression in iPSC-PPC. Box plots colored in black indicate genes with a significant $e_gQTL$, while the one colored in gray indicates that it was not associated with genotype. Box plots indicate median interquartile range (IQR), and 1.5 x IQR. Smoothed regression line represents the relationship between genotype and normalized expression (*lm* function in R). We note that *ST13* is an eGene in iPSC-PPC but its $e_gQTL$ signal neither colocalized nor was in LD with *ST13* $e_gQTLs$ in adult islet and adult whole pancreas. The signal was significant (q-value < 0.01) but weak.

#### Supplementary Figure 15: Pancreatic $e_g$QTL colocalization with GWAS traits

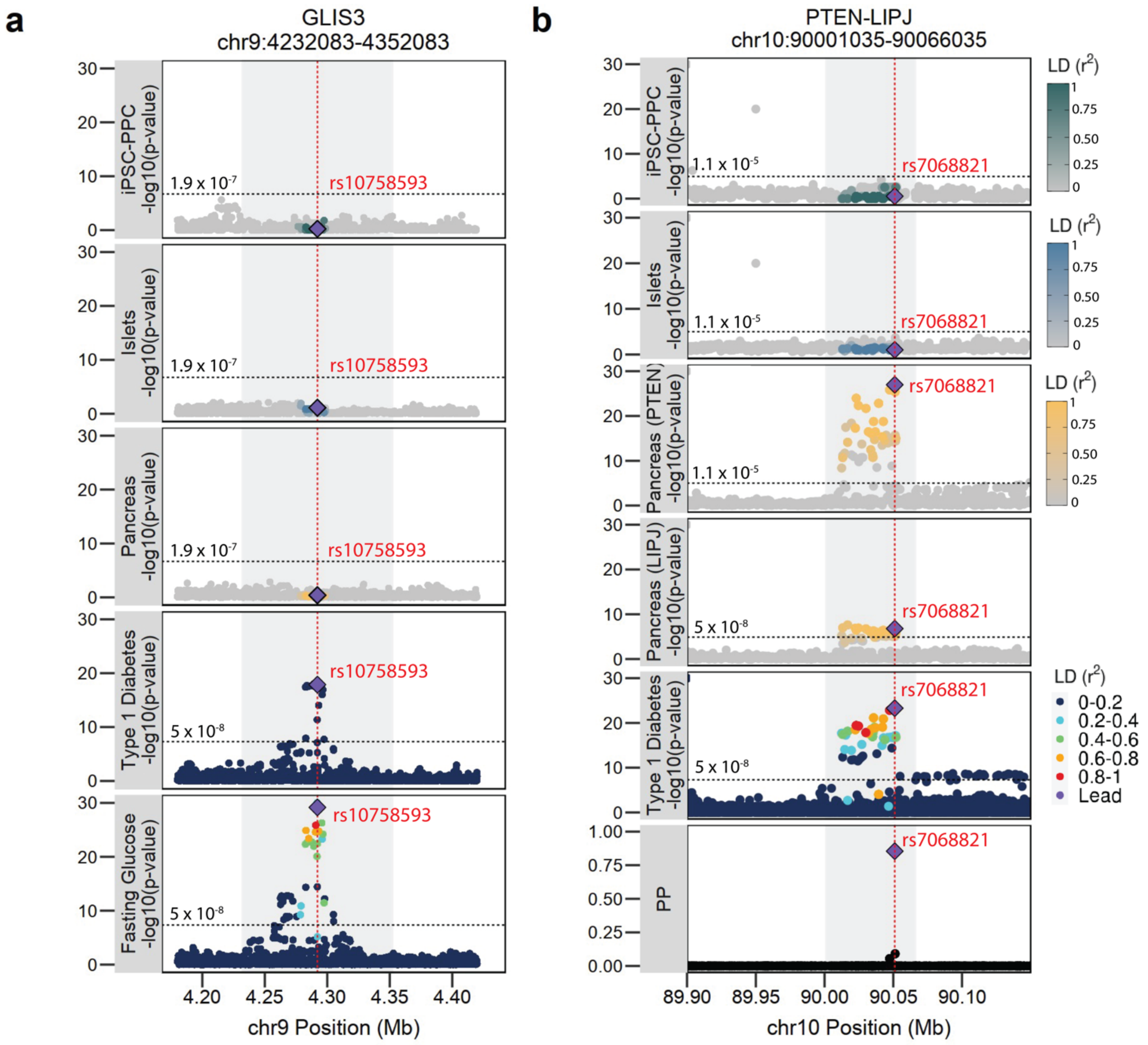


(**a**) This panel shows that the GWAS signals associated with T1D-risk and FG levels in the chr9:423083-4352083 locus did not colocalize with $e_g$QTLs for *GLIS3* in iPSC-PPC and the adult pancreatic tissues. (**b**) The GWAS signal associated with T1D-risk in the chr10:90001035-90066035 locus colocalized with an adult whole pancreas-unique $e_g$QTL module containing $e_g$QTLs for *PTEN* and *LIPJ*. Each point in these plots represents an individual SNP color-coded by LD with the lead candidate causal variant highlighted in purple. The bottom plot in panel **b** shows the posterior probability (PP) of association for each variant being causal for both $e_g$QTL and GWAS associations. For plotting purposes, we assigned a single p-value for gene-level significance based on Bonferroni-correction (0.05 divided by the number of variants tested for the gene; horizontal line) for the $e_g$QTL signals. For iPSC-PPC and adult islet in panel **b**, we overlaid eQTL associations for *PTEN, LIPJ*, and nearby genes to show that the locus was not associated with gene expression in the tissues. For GWAS signals, we used p-value = $5x10^{-8}$ to indicate genome-wide significance. Red vertical lines indicate the positions of the lead candidate causal variants underlying GWAS and eQTL colocalization based on maximum PP.

#### Supplementary Figure 16: e$_{AS}$QTL associations for GWAS-associated eIsoforms in fetal-like iPSC-PPC

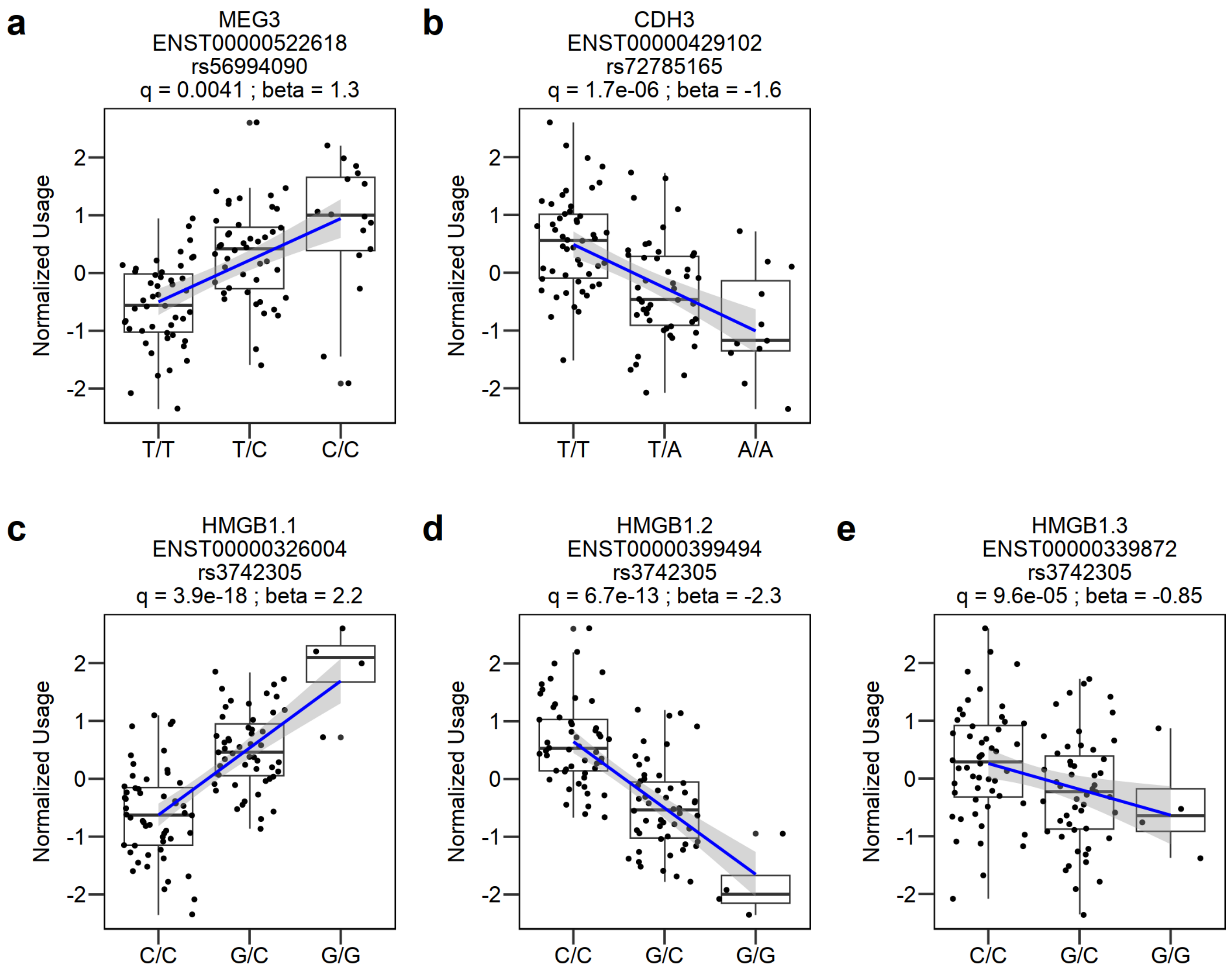


For each candidate susceptibility isoform, we show the association between the genotype of the lead candidate causal variant (predicted from GWAS-eQTL colocalization) and their normalized isoform usage in iPSC-PPC. Box plots indicate median interquartile range (IQR), and 1.5 x IQR. Smoothed regression line indicate the relationship between genotype and normalized expression (*lm* function in R).

#### Supplementary Figure 17: PEER Factor Optimization

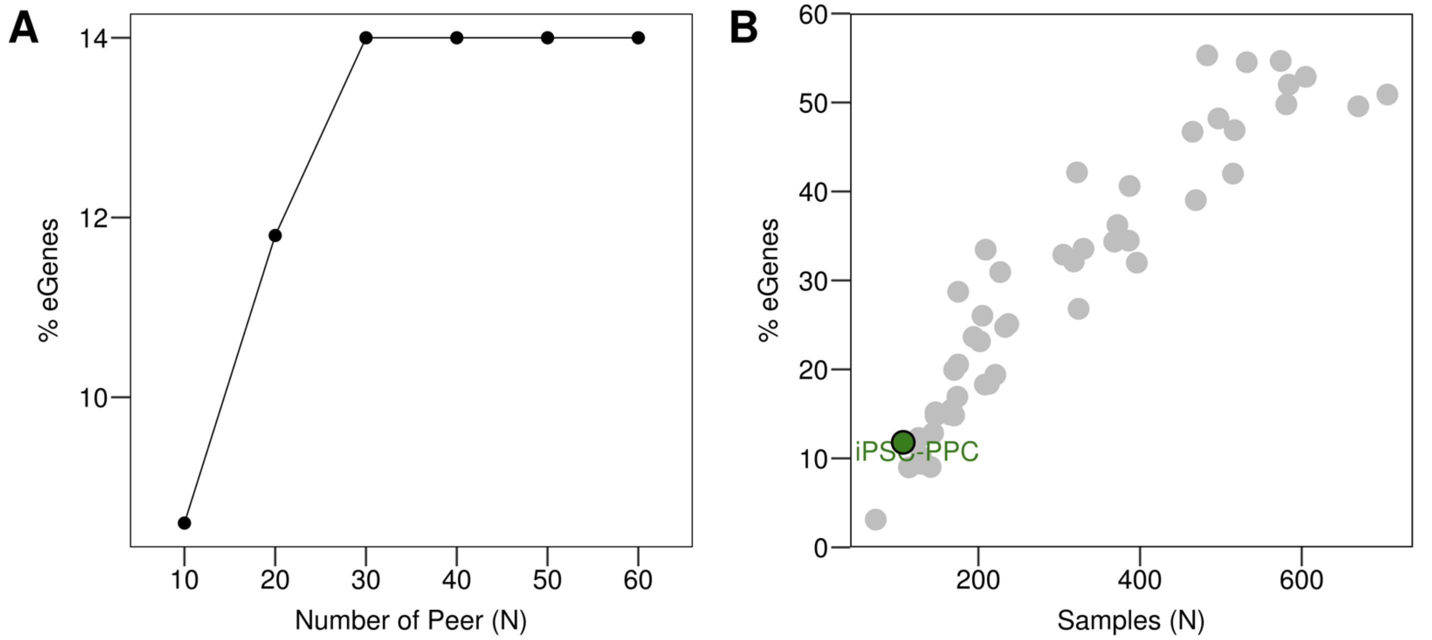


The line plot in panel **a** shows the percentage of eGenes discovered in iPSC-PPC on a random set of 500 genes using PEER factors ranging from 10 to 60 in increments of 10 as covariates. While 30 PEER factors resulted in maximum eGene percentage (14.0%), we selected 20 PEER factors (11.8%) because the percentage of eGenes was comparable to GTEx tissues of similar sample sizes (panel **b**; colored in gray). These results show that compared with GTEx, our study was as well-powered to detect significant eQTL associations.

#### Supplementary Figure 18: Correlation between eQTL covariates

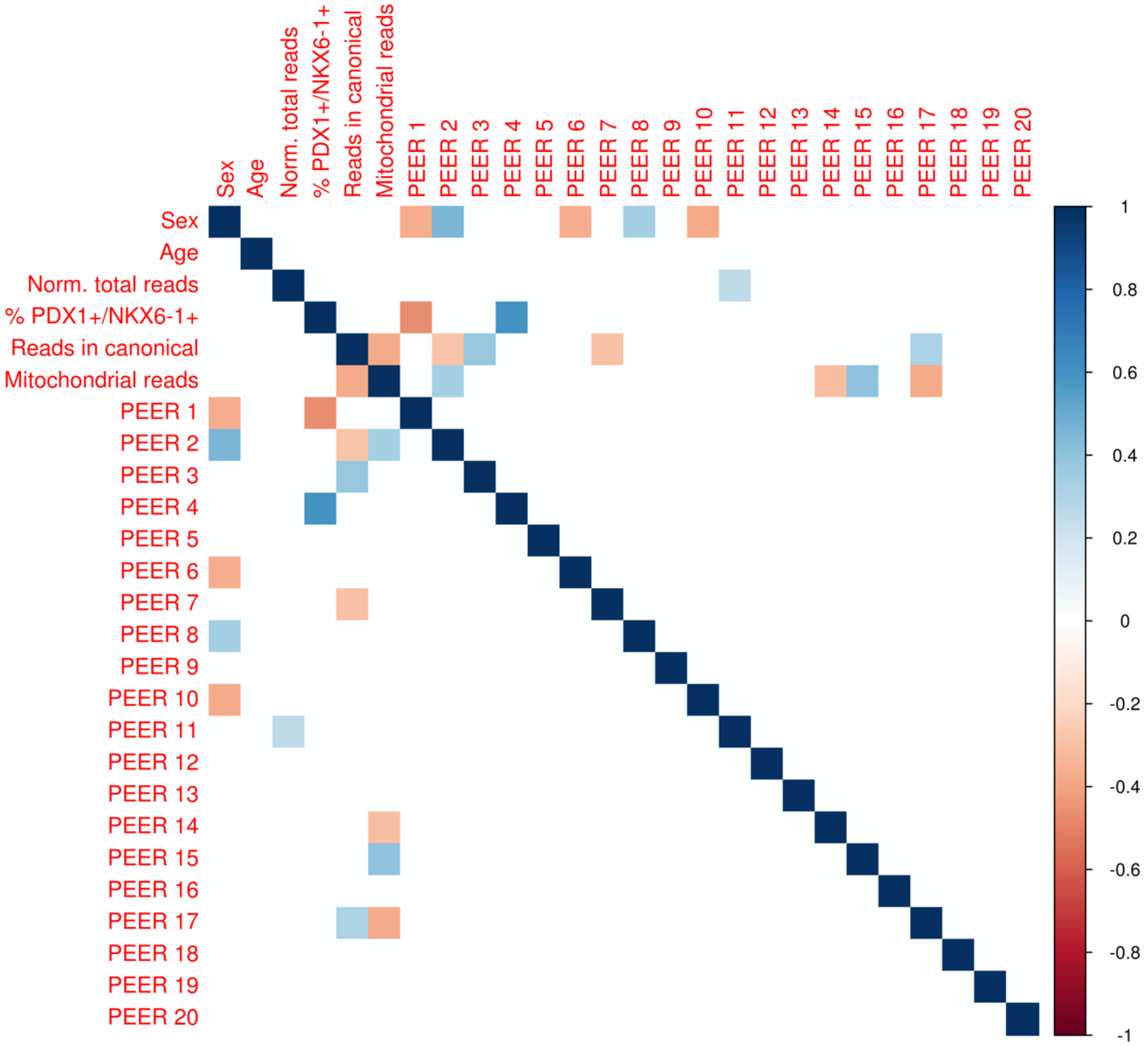


To determine whether PEER factors were correlated with other attributes of iPSC-PPC, we performed a spearman correlation analysis between each pair of covariates used in the linear mixed model for eQTL analysis. We found that PEER factors 1 and 4 were associated with the percentage of cells expressing PDX1$^{+}$/NKX6-1$^{+}$ based on FACs, therefore accounting for any cellular heterogeneity in iPSC-PPC that could be driving expression variability across the samples.

#### Supplementary Data Legends

##### Supplementary Data 1: Subject Information

The first sheet contains information about each of the 106 iPSCORE individuals included in this study. Columns are: **ipscore_id**: subject iPSCORE ID, **subject_uuid**: universally unique identifier (UUID) for the subject, **sex**: sex, **age**: age at the time of enrollment, **most_similar_1kg_pop**: most similar 1000 Genomes Phase 3 superpopulation [9], **family_uuid**: family UUID, **wgs_uuid**: whole-genome sequencing (WGS) sample UUID, and **pc1-20**: the first 20 genotype principal components (PC) accounting for global ancestry. The second sheet contains the kinship matrix used for eQTL mapping, where the row and column names correspond to the WGS UUID's.

##### Supplementary Data 2: Sample Information

This table contains information about the 107 iPSC-PPC samples used in this study. Columns are: **udid**: unique differentiation ID, **subject_uuid**: subject UUID, **wgs_uuid**: WGS sample UUID, **live_scrna_uuid**: sample UUIDs for scRNA-seq samples prepared with fresh cells, **cryo_scrna_pool_uuid**: sample UUIDs for scRNA-seq samples prepared after cryopreservation of the cells (cells from four iPSC-PPC samples were pooled into one sample and cells from another three iPSC-PPC samples were pooled into a second sample), **cryo_scrna_pool_name**: name labels for the pooled scRNA-seq samples, **live_scrna_pi_hat**: PI_HAT indicating sample match to the subject (from plink genome), **bulk_rna_uuid**: sample UUIDs for each bulk RNA-seq sample, **ipsc_clone**: clone of the iPSC line used for differentiation, **ipsc_passage_at_monolayer**: passage of the iPSC line at monolayer, **day15_pdx1_nkx6.1**: percentage of cells expressing PDX1 and NKX6-1 at day 15 of differentiation measured by flow cytometry, **day15_pdx1**: percentage of cells expressing PDX1 at day 15 of differentiation measured by flow cytometry, **day15_nkx6.1**: percentage of cells expressing NKX6-1 at day 15 of differentiation measured by flow cytometry, **total_reads**: total reads sequenced (note: divide this number by 2 to get the number of paired reads), **total_reads_norm**: normalized total number of reads sequenced, **uniquely_mapped_reads_canonical_chromosomes**: percentage of uniquely mapped reads in autosomal and sex chromosomes, **pct_intergenic_bases**: percentage of bases that mapped to intergenic regions of genomic DNA (from Picard RnaSeqMetrics), **pct_mrna_bases**: percentage of bases that mapped to regions corresponding to UTRs and coding regions of mRNA transcripts (from Picard RnaSeqMetrics), **pct_duplicates**: percentage of duplicate reads (from samtools flagstat), **pct_mitochondrial_reads**: percentage of reads mapping to mitochondrial chromosome (from samtools idxstats), **bulk_rna_pi_hat**: PI_HAT indicating sample match to the subject (from plink genome), **peer1-20**: the 20 PEER factors used in eQTL mapping in iPSC-PPC.

##### Supplementary Data 3: scRNA-seq metadata

For each of the 84,225 single cells that passed quality control, we provide: **cell_id**: ID of the single cell, **barcode:** cell barcode, **cryo_scrna_pool_name:** name labels for the pooled scRNA-seq samples (only for samples prepared using cryopreserved cells), **sample_preparation**: method of scRNA-seq preparation ("fresh" indicates that the scRNA-seq sample was prepared using fresh cells immediately after differentiation, "Cryopreserved" indicates that the scRNA-seq sample was prepared using cryopreserved cells), **udid**: unique differentiation ID, **scrna_uuid**: UUID for the scRNA-seq sample the cells came from (corresponds to **live_scrna_uuid** and **cryo_scrna_pool_uuid** columns in Supplementary Data 2), **wgs_uuid**: WGS sample UUID, **subject_uuid**: subject UUID, **ncount_rna**: total number of molecules detected within a cell, **nfeature_rna**: total number of genes detected in each cell, **percent_mt**: proportion of transcripts mapping to mitochondrial genes, **cluster_res.0.05**: cluster ID numbers for each cell at resolution 0.05, **cluster_res.0.08**: cluster ID numbers for each cell at resolution 0.08, **cluster_res.0.1**: cluster ID numbers for each cell at resolution 0.1, **celltype**: cell type labels for each cell at resolution 0.08, **UMAP_1** and **UMAP_2**: UMAP coordinates of each cell. The WGS UUIDs were mapped to each single cell using Demuxlet [22] and then mapped to the subject UUID. A Seurat R Object for filtered and integrated dataset is available on Figshare: https://doi.org/10.6084/m9.figshare.21836208.

**Supplementary Data 4: Differentially expressed genes in scRNA-seq clusters**

For each celltype and gene, we report: **pct.1**: percentage of cells that expressed the gene in the cell type cluster, **pct.2**: percentage of cells that expressed the gene outside of the celltype cluster, **avg_log2FC:** the log fold-change of the average expression between the two groups, **p_val**: p-value from two-sided Wilcox Rank-Sum test, **p_val_adj**: adjusted p-value based on Bonferroni correction using all features in the dataset. Genes with adjusted p-value ≤ 0.05 were considered differentially expressed.

**Supplementary Data 5: Cellular deconvolution of iPSC-PPC bulk RNA-seq**

This table reports the estimated relative proportions of each of the eight cell types (clusters) identified in the scRNA-seq data for each of the 107 iPSC-PPC bulk RNA-seq samples (Supplementary Figure 4). Columns are: **UDID**: unique differentiation ID, **Early_DE**: the estimated relative proportion of early DE cells, **Early_Ductal**: the estimated relative proportion of early ductal cells, **Early_PPC**: the estimated relative proportion of early PPC cells, **Endocrine**: the estimated relative proportion of endocrine cells, **iPSC**: the estimate relative proportion of iPSC cells, **Late_PPC**: the estimated relative proportion of late PPC cells, **Mesendoderm**: the estimated relative proportion of mesendoderm cells, **Rep_Late_PPC**: the estimated relative proportion of replicating late PPC cells. The relative proportions were estimated by CIBERSORTx [10] deconvolution (Supplementary Figure 7B). In sheet 2, we provide the cell type signature matrix used for the deconvolution.

**Supplementary Data 6: PCA and pseudotime analyses on iPSC, iPSC-PPC, and adult pancreatic tissues**

We report results from PCA and pseudotime analysis on 107 iPSC-PPCs, 213 iPSCs [11], 87 adult islets [12], and 176 adult whole pancreas samples (phs000424.v7.p2) (Supplementary Figure 9). For each sample, we report: **sample_id**: bulk RNA-seq ID, **tissue**: tissue type, **pseudotime**: pseudotime inferred by Monocle [14] (see Methods), **pc1-10:** principal components for each sample after principal components analysis on the top variable genes (see Methods). Sample IDs were assigned as the following: sample bulk RNA-seq UUIDs for iPSCORE samples (iPSC and iPSC-PPC), sample accession IDs from GEO for adult islet samples (GSE50398), and SRA accession IDs for adult whole pancreas samples (phs000424.v7).

**Supplementary Data 7: Lead variants for all $e_g$QTLs and $e_i$QTLs in iPSC-PPC**

The table reports the lead eSNP for each eQTL discovered in iPSC-PPC. We provide: **eqtl_phenotype**: the phenotype the eQTL was associated with (gene expression or isoform usage), **transcript_id**: transcript ID, **gene_id:** gene ID, **gene_name**: gene name, **discovery_order**: discovery order of the eQTL, where 0 represents primary eQTL and 1-4 represents conditional eQTLs, **eqtl_id**: eQTL ID assigned as [tissue_type]_[discovery_order]_[transcript_ID], **snp**: lead variant ID assigned as [chromosome]_[ position]_[reference allele]_[alternate allele], **chrom**: the lead variant's chromosome, **pos:** hg19 position of the lead variant, **ref**: reference allele of the lead variant, **alt**: alternate allele of the lead variant, **beta**: the lead variant's effect size on gene expression or isoform usage, **se**: standard error, **pval**: eQTL p-value for the association between genotype of the lead variant and gene expression or isoform usage, **tests**: number of independent variants used for eigenMT [23] p-value correction, **fdr**: FDR-corrected eQTL p-value calculated by eigenMT, **qval**: q-value from Benjamini-Hochberg correction of fdr, **egene:** TRUE/FALSE indicating whether the eQTL is significant or not with q-value threshold < 0.01. Full summary statistics are available on Figshare as a tar zipped directory containing text files for each gene and isoform tested: https://doi.org/10.6084/m9.figshare.21899496 and https://doi.org/10.6084/m9.figshare.21899499.

**Supplementary Data 8: iPSC-PPC PPC eQTL Credible Sets of SNPs with PP ≥ 1%**

For each significant iPSC-PPC $e_g$QTL and $e_i$QTL, we report their fine-mapped variants with causal PP ≥ 1%. eQTLs that are missing from this table do not have SNPs above the PP threshold. We provide: **eqtl_phenotype**: the molecular phenotype tested for association with genetic variation (gene expression or isoform usage), **eqtl_id**: eQTL ID assigned as [tissue_type]_[discovery_order]_[transcript_ID], **transcript_id**: transcript ID, **gene_id**: gene ID, **discovery_order**: discovery order of the eQTL, where 0 represents primary eQTL and 1-4 represents conditional eQTLs, **snp**: lead variant ID assigned as [chromosome]_[ position]_[reference allele]_[alternate allele], fine-mapping statistics from *finemap.abf* (columns G-N) in *coloc* [24], and **SNP.PP**: posterior probability that the variant is causal for the association between genotype and gene expression or isoform usage. Tables for $e_g$QTLs (key name: "gene") and $e_i$QTLs (key name: "isoform") are available on Figshare as an R list object due to large sample size: https://doi.org/10.6084/m9.figshare.21836211.

**Supplementary Data 9: Individual SNP TF Binding Impact Score using GVAT Database**

We report GVATdb [25] results for each eQTL using their fine-mapped variants with causal PP ≥ 10%. The table provides: **eqtl_id**: eQTL ID assigned as [tissue_type]_[discovery_order]_[transcript_ID], **snp_beta**: the variant's effect size on gene expression or isoform usage from eQTL analysis, **SNP.PP**: posterior probability that the variant is causal for the association between genotype and gene expression or isoform usage, **tf**: name of the transcription factor tested for binding effects, **allele1_bind**: TF binding score for reference allele, **allele2_bind**: TF binding score for alternate allele, **seq_binding**: label yes (Y) or no (N) indicating whether the TF binds to the variant, **deltaSVM_score**: impact score of TF binding by the variant, **preferred_allele**: label indicating whether the variant results in "Gain", "Loss", or has no effects ("None") on TF binding. Tables for $e_g$QTLs (key name: "gene") and $e_i$QTLs (key name: "isoform") are available on Figshare as an R list object due to large file size: https://doi.org/10.6084/m9.figshare.21836211.

**Supplementary Data 10: Correlation between eQTL effect sizes and TF binding affinity**

This table contains results for the association analysis between eQTLs and TF binding affinity. We report: **snp.pp_threshold:** threshold used for individual variant causal posterior probability, **eqtl_phenotype**: the molecular phenotype tested for association with genetic variation (gene expression or isoform usage), **cor**: estimated measure of association from Pearson's product-moment correlation, **pval**: p-value of the correlation test.

**Supplementary Data 11: Colocalization Results between iPSC-PPC and Adult eQTLs (PP ≥ 80%)**

Sheet 1: Colocalization between iPSC-PPC egQTL and eiQTLs. The table reports colocalization results for the 410 eGenes with H3 and/or H4 associations between their $e_g$QTLs and corresponding $e_i$QTLs.

Sheet 2: eGene colocalization between iPSC-PPC and adult islet. The table reports colocalization results for the 795 shared eGenes with H3 and/or H4 association between iPSC-PPC and adult islet $e_g$QTLs.

Sheet 3: Input for generating $e_g$QTL networks. The table reports colocalization results for all 7,893 $e_g$QTL pairs between the three pancreatic tissues.

Sheet 4: Input for generating $e_{AS}$QTL networks. The table reports colocalization results for all 4,868 $e_{AS}$QTLs pairs between the three pancreatic tissues.

In each table, we provide: **eqtl_id.1**: eQTL ID for one of the two eQTLs being colocalized, **eqtl_id.2**: eQTL ID for thw second eQTL being colocalized, **transcript_id.1**: transcript ID for eqtl_id.1, **transcript_id.2**: transcript ID for eqtl_id.2, **gene_id.1**: gene ID for eqtl_id.1, **gene_id.2**: gene ID for eqtl_id.2, **gene_name.1**: gene name for eqtl_id.1, **gene_name.2**: gene name for eqtl_id.2, **eqtl_phenotype.1**: the molecular phenotype tested for association with genetic variation (gene expression or isoform usage) for **eqtl_id.1**, **eqtl_phenotype.2**: the molecular phenotype tested for association with genetic variation (gene expression or isoform usage) for **eqtl_id.2**, **tissue.1**: tissue the first eQTL was detected in, **tissue.2**: the tissue the second eQTL was detected in, **discovery_order**: discovery order of the eQTL, where 0 represents primary eQTL and 1-4 represents conditional eQTLs, **nsnps**: number of variants used to test for colocalization (obtained from *coloc.abf*), **PP.H0.abf**: posterior probability of H0 model (no causal variant), **PP.H1.abf**: posterior probability of H1 model (causal variant for trait 1 only), **PP.H2.abf**: posterior probability of H2 model (causal variant for trait 2 only), **PP.H3.abf**: posterior probability of H3 model (two distinct causal variants), **PP.H4.abf**: posterior probability of H4 model (one common causal variant), **likely_model**: model with the strongest evidence of being true based on highest posterior probability, **max_model_pp**: the maximum PP across the models, **topsnp**: the lead predicted causal variant if PP.H4.abf was true, **topsnp_pp**: the posterior probability that **topsnp** is causal for the association with the molecular phenotype. Multiple variants may be listed as the lead predicted causal variants if they share the same maximal posterior probability. eQTL IDs were assigned as [tissue_type]_[discovery_order]_[transcript_ID].

**Supplementary Data 12: eQTL Annotation for each iPSC-PPC and Adult eQTLs**

The table describes the annotations for each $e_g$QTL and $e_{AS}$QTL in the three pancreatic tissues. We provide: **eqtl_id:** eQTL ID assigned as [tissue_type]_[discovery_order]_[transcript_ID], **transcript_id:** transcript ID, **gene_id**: gene ID, **gene_name**: gene_name, **tissue**: tissue source the eQTL was detected in, **eqtl_phenotype**: the molecular phenotype tested for association with genetic variation (gene expression or alternative splicing), **eqtl_type**: label indicating whether the eQTL was a singleton or combinatorial, **module_id**: module ID assigned as [eQTL_phenotype]_[chromosome]_[number], where "GE" represents eQTL modules associated withe gene expression and "AS" represents eQTL modules associated with alternative splicing (module IDs are given to only combinatorial eQTLs, see also Supplementary Data 13), **expressed_ipsc_ppc**: TRUE/FALSE indicating whether the gene was expressed and tested for genetic association in iPSC, **expressed_islet**: TRUE/FALSE indicating whether the gene was expressed and tested for genetic association in adult islets, **expressed_pancreas**: TRUE/FALSE indicating whether the gene was expressed and tested for genetic association in adult whole pancreas, **LD_ipsc_ppc:** TRUE/FALSE indicating whether the eQTL was in LD with nearby iPSC-PPC eQTLs, **LD_islet**: TRUE/FALSE indicating whether the eQTL was in LD with nearby adult islet eQTLs, **LD_pancreas**: TRUE/FALSE indicating whether the eQTL was in LD with nearby adult whole pancreas eQTLs, **islet_egene_overlap**: labels describing the eGene overlap between iPSC-PPC and adult islet eQTLs in the module (zero means there were no adult islet eQTLs in the module, same means that all eGenes overlapped between iPSC-PPC and adult islet eQTLs, partial means that there was at least one shared eGene and at least one different eGene between iPSC-PPC and adult islet eQTLs, and different means that there was no overlap in eGenes between iPSC-PPC and adult islet eQTLs), **pancreas_egene_overlap**: labels describing the eGene overlap between iPSC-PPC and adult whole pancreas eQTLs in the module (zero means there were no adult whole pancreas eQTLs in the module, same means that all eGenes were the same between all iPSC-PPC and adult whole pancreas eQTLs, partial means that there was at least one shared eGene and at least one different eGene between iPSC-PPC and adult whole pancreas eQTLs, and different means that there was no overlap in eGenes between iPSC-PPC and adult whole pancreas eQTLs), **module_pass**: TRUE/FALSE indicating whether the module passed threshold requirements (see Methods), **category_annotation**: labels for for each eQTL based on whether it was unique to a single tissue, shared with another tissue, or was a singleton or combinatorial (see below or Methods for descriptions for each category), **notes**: comments describing why the eQTL was annotated as "ambiguous" or "module_failed". Below, we describe what each category annotation means in the table. Descriptions are also provided in the Methods.

1) "ipsc_ppc singleton": the eQTL was an iPSC-PPC-unique singleton eQTL
2) "islet singleton": the eQTL was an adult islet-unique singleton eQTL
3) "whole-pancreas singleton": the eQTL was an adult whole pancreas-unique singleton eQTL
4) "ipsc_ppc-unique": the eQTL was in an iPSC-PPC-unique module
5) "islet-unique": the eQTL was in an adult islet-unique module
6) "whole-pancreas-unique": the eQTL was in an adult whole pancreas-unique module
7) "adult-shared": the eQTL was in an adult-shared module (shared between adult islets and adult whole pancreas; module contained at least one eQTL from adult islets, at least one eQTL from adult whole pancreas, and zero eQTLs from iPSC-PPC)
8) "fetal-islet": the eQTL was in a fetal-islet module (shared between iPSC-PPC and adult islets; module contained at least one eQTL from iPSC-PPC, at least one eQTL from adult islets, and zero eQTLs from adult whole pancreas)

9) "fetal-whole-pancreas": the eQTL was in a fetal-whole-pancreas module (shared between iPSC-PPC and adult whole pancreas; module contained at least one eQTL from iPSC-PPC, at least one eQTL from adult whole pancreas, and zero eQTLs from adult islets)
10) "fetal-adult": the eQTL was in a fetal-adult module (shared between iPSC-PPC and the two adult tissues; module contained at least one eQTL from iPSC-PPC, at least one eQTL from adult islets, and at least one eQTL from adult whole pancreas)
11) "module_failed": the eQTL was excluded due to being in a module that did not satisfy threshold requirements (see Methods)
12) "ambiguous": the eQTL was excluded due to being in LD with a nearby eQTL. If the eQTL was in a module, the eQTL was in LD with an eQTL in a different tissue. If the eQTL was a singleton eQTL, the eQTL was in LD with another nearby eQTL in the same or different tissue. Tissue-specificity for this eQTL could not be determined and therefore excluded from downstream analyses.

**Supplementary Data 13: Network Modules of iPSC-PPC and Adult eQTLs**

The table provides information for each $e_g$QTL (sheet 1) and $e_{AS}$QTL (sheet 2) module. Specifically, we provide: **module_id**: module ID assigned as [eQTL_phenotype]_[chromosome]_[number], where "GE" represents eQTL modules associated withe gene expression and "AS" represents eQTL modules associated with alternative splicing (module IDs are given to only combinatorial eQTLs), **associations:** all eQTL associations in the module by their eQTL IDs, **number_assocs:** number of eQTL associations in the module, **number_ipsc_ppc_assocs**: number of iPSC-PPC eQTL associations in the module, **number_islet_assocs**: number of adult islet eQTL associations in the module, **number_pancreas_assocs:** number of adult whole pancreas eQTL assocations in the module, **islet_egene_overlap**: labels describing the eGene overlap between iPSC-PPC and adult islet eQTLs in the module (zero means there were no adult islet eQTLs in the module, same means that all eGenes overlapped between iPSC-PPC and adult islet eQTLs, partial means that there was at least one shared eGene and at least one different eGene between iPSC-PPC and adult islet eQTLs, and different means that there was no overlap in eGenes between iPSC-PPC and adult islet eQTLs), **pancreas_egene_overlap**: labels describing the eGene overlap between iPSC-PPC and adult whole pancreas eQTLs in the module (zero means there were no adult whole pancreas eQTLs in the module, same means that all eGenes were the same between all iPSC-PPC and adult whole pancreas eQTLs, partial means that there was at least one shared eGene and at least one different eGene between iPSC-PPC and adult whole pancreas eQTLs, and different means that there was no overlap in eGenes between iPSC-PPC and adult whole pancreas eQTLs), **egene_overlap_category**: eGene overlap category shown in Figure 4A and Supplementary Figure 11C, **module_pass**: TRUE/FALSE indicating whether the module passed threshold requirements (see Methods), **category_annotation**: labels for for each eQTL based on whether it was unique to a single tissue, shared with another tissue, or was a singleton or combinatorial (see below or Methods for descriptions for each category), **notes**: comments describing why the eQTL was annotated as "ambiguous" or "module_failed". Below, we describe what each category annotation means in this table. Descriptions are also provided in the Methods.

1) "ipsc_ppc-unique": the module contained only iPSC-PPC eQTLs
2) "islet-unique": the module contained only adult islet eQTLs

3) “whole-pancreas-unique”: the module contained only adult whole pancreas eQTLs
4) “adult-shared”: the module was an adult-shared module and contained only eQTLs from adult islet and adult whole pancreas
5) “fetal-islet”: the module contained only iPSC-PPC and adult islet eQTLs
6) “fetal-whole-pancreas”: the module contained only iPSC_PPC and adult whole pancreas eQTLs
7) “fetal-adult”: the module contained iPSC-PPC, adult islet, and adult whole pancreas eQTLs
8) “module_failed”: the module did not pass threshold requirements (see Methods)
9) “ambiguous”: the module contained an eQTL that is in LD with another eQTL in a different tissue

**Supplementary Data 14: Chromatin State Enrichments of iPSC-PPC-unique eQTL Singletons and Modules**

The table reports chromatin enrichment results for iPSC-PPC-unique singleton and combinatorial $e_g$QTLs at various SNP.PP thresholds. We provide: **snp.pp_threshold:** threshold used for individual variant causal posterior probability, **category_annotation**: labels for for each eQTL based on whether it was unique to a single tissue, shared with another tissue, or was a singleton or combinatorial (see above or Methods for descriptions for each category), **eqtl_type**: label indicating whether the eQTL was a singleton or combinatorial, **chromatin_annotation**: chromatin state annotation, **tissue**: the tissue where the chromatin annotations were drived from, **study**: the study source for the chromosome annotation, **estimate**: odds-ratio, **pval**: p-value of significance for two-sided Fisher’s Exact Test comparing the fraction of variants overlapping the chromotation annotations against a null set of variants, **qval:** Benjamini-Hochberg correction of the p-value. **ci1** and **ci2**: lower and upper limits of the confidence interval.

**Supplementary Data 15: Colocalization of iPSC-PPC and Adult eQTLs with Pancreatic GWAS Traits**

The table reports information about the 312 GWAS loci that colocalized with iPSC-PPC and/or adult eQTLs with PP.H4 ≥ 80%. Note that a GWAS locus can be listed more than once if it colocalized with more than one eQTL (only observed for modules). In this table, we provide: **gwas_locus_id**: eQTL-GWAS locus ID assigned as [eQTL module ID or eQTL singleton ID] [trait_id], **eqtl_id**: eQTL ID assigned as [tissue_type]_[discovery_order]_[transcript_ID], **transcript_id**: transcript ID**, gene_id**: gene ID, **gene_name**: gene name, **eqtl_phenotype**: the molecular phenotype tested for association with genetic variation (gene expression or alternative splicing), **tissue**: tissue source the eQTL was detected in, **eqtl_type**: label indicating whether the eQTL was a singleton or combinatorial, **module_id**: module ID assigned as [eQTL_phenotype]_[chromosome]_[number], where “GE” represents eQTL modules associated withe gene expression and “AS” represents eQTL modules associated with alternative splicing (module IDs are given to only combinatorial eQTLs, see also Supplementary Data 13), **trait_id**: ID for the GWAS trait, **description**: description of the GWAS trait, **study_source**: study source of the GWAS summary statistics, **nsnps**: number of variants used to test for colocalization between GWAS and eQTL (obtained from *coloc.abf*), **PP.H0.abf**: posterior probability of H0 model (no causal variant), **PP.H1.abf**: posterior probability of H1 model (causal variant for trait 1 only), **PP.H2.abf**: posterior probability of H2 model (causal variant for trait 2 only), **PP.H3.abf**: posterior probability of H3 model (two distinct causal variants), **PP.H4.abf**: posterior probability of H4 model (one common causal variant), **max_model_pp**: the maximum PP across the models, **likely_model**: model with the strongest evidence of being true based on highest posterior probability, **topsnp**: the lead predicted causal variant if PP.H4.abf was true, **topsnp_pp**: the posterior probability that **topsnp** is causal for the association with the molecular phenotype, **topsnp_gwas_pval**: GWAS p-value for the association between the GWAS trait and the variant, **topsnp_eqtl_pval**: eQTL p-value for the association between gene expression/alternative splicing and the variant, **islet_egene_overlap**: labels describing the eGene overlap between iPSC-PPC and adult islet eQTLs in the module (zero means there were no adult islet eQTLs in the module, same means that all eGenes overlapped between iPSC-PPC and adult islet eQTLs, partial means that there was at least one shared eGene and at least one different eGene between iPSC-PPC and adult islet eQTLs, and different means that there was no overlap in eGenes between iPSC-PPC and adult islet eQTLs), **pancreas_egene_overlap**: labels describing the eGene overlap between iPSC-PPC and adult whole pancreas eQTLs in the module (zero means there were no adult whole pancreas eQTLs in the module, same means that all eGenes were the same between all iPSC-PPC and adult whole pancreas eQTLs, partial means that there was at least one shared eGene and at least one different eGene between iPSC-PPC and adult whole pancreas eQTLs, and different means that there was no overlap in eGenes between iPSC-PPC and adult whole pancreas eQTLs), **category_annotation**: labels for for each eQTL based on whether it was unique to a single tissue, shared with another tissue, or was a singleton or combinatorial (see Supplementary Data 12 and 13 for the description of each category), **used_to_finemap**: the eQTL used to finemap the GWAS signal (relevant for eQTL modules that had multiple eQTL signals that colocalized with the GWAS signal), **cs_size**: number of putative causal variants in the 99% credible set. Multiple variants may be listed as the lead predicted causal variants (**topsnp**) if they share the same maximal posterior probability

**Supplementary Data 16: 99% Credible Sets for iPSC-PPC eQTLs Associated with GWAS**

This table contains the 99% credible sets for each of the 312 GWAS loci described in Supplementary Data 15. The eQTLs used to construct the 99% credible sets can be found in Supplementary Data 15 with the **used_to_finemap** column. In this table, we provide: **gwas_locus_id**: eQTL-GWAS locus ID assigned as [eQTL module ID or eQTL singleton ID] [trait_id], **trait_id**: ID for the GWAS trait, **description**: description of the GWAS trait, **study_source**: study source of the GWAS summary statistics, **eqtl_id**: eQTL ID assigned as [tissue_type]_[discovery_order]_[transcript_ID], **snp**: ID for the variant in the credible set, colocalization statistics from *coloc.abf* (columns F-T), **SNP.PP.H4**: posterior probability for the variant being causal for eQTL and GWAS associations from *coloc.abf*.

**Supplementary Data 17: LD analysis with non-pancreatic GTEx tissues**

This table contains results for the LD analysis conducted between the 16 iPSC-PPC-unique $e_g$QTLs that colocalized with GWAS signals and the $e_g$QTLs in the 48 non-pancreatic tissues in the GTEx dataset version 8 [4]. We provide: **eqtl_id:** eQTL ID assigned as [tissue_type]_[discovery_order]_[transcript_ID], **eqtl_phenotype**: the molecular phenotype tested for association with genetic variation (gene expression or alternative splicing), **eqtl_type**: label indicating whether the eQTL was a singleton or combinatorial, **tissue**: tissue source the eQTL was detected in, **transcript_id**: transcript ID, **gene_id**: gene ID, **gene_name**: gene name, **category_annotation**: labels for for each eQTL based on whether it was unique to a single tissue, shared with another tissue, or was a singleton or combinatorial (see Supplementary Data 12 and 13 for the description of each category), **in_ld_other_gtex**: TRUE/FALSE indicating whether the $e_gQTL$ is in LD with an adult adult $e_gQTL$
